# High-content screening of coronavirus genes for innate immune suppression reveals enhanced potency of SARS-CoV-2 proteins

**DOI:** 10.1101/2021.03.02.433434

**Authors:** Erika J. Olson, David M. Brown, Timothy Z. Chang, Lin Ding, Tai L. Ng, H. Sloane Weiss, Yukiye Koide, Peter Koch, Nathan Rollins, Pia Mach, Tobias Meisinger, Trenton Bricken, Joshua Rollins, Yun Zhang, Colin Molloy, Yun Zhang, Bridget N. Queenan, Timothy Mitchison, Debora Marks, Jeffrey C. Way, John I. Glass, Pamela A. Silver

## Abstract

Suppression of the host intracellular innate immune system is an essential aspect of viral replication. Here, we developed a suite of medium-throughput high-content cell-based assays to reveal the effect of individual coronavirus proteins on antiviral innate immune pathways. Using these assays, we screened the 196 protein products of seven coronaviruses (SARS-CoV-2, SARS-CoV-1, 229E, NL63, OC43, HKU1 and MERS). This includes a previously unidentified gene in SARS-CoV-2 encoded within the Spike gene. We observe immune-suppressing activity in both known host-suppressing genes (e.g., NSP1, Orf6, NSP3, and NSP5) as well as other coronavirus genes, including the newly identified SARS-CoV-2 protein. Moreover, the genes encoded by SARS-CoV-2 are generally more potent immune suppressors than their homologues from the other coronaviruses. This suite of pathway-based and mechanism-agnostic assays could serve as the basis for rapid *in vitro* prediction of the pathogenicity of novel viruses based on provision of sequence information alone.

## Introduction

Innate immunity is the human body’s first line of defense against pathogens. We and others hypothesize that viral pathogenicity may correlate with the degree to which a virus inhibits the antiviral innate immune response. For example, Ebola virus encodes several genes that suppress innate immune signaling, with the result that this virus can reach extremely high titers before the immune system mounts an effective response (Kuhl et al., 2012). Hoffmann et al. (2015) described the interaction between the host innate immune response and its suppression by viral genes as a ‘molecular arms race’ and summarized how most or all viruses encode such suppressing factors. Mechanisms of suppression include: inhibition of nuclear protein transport, proteolysis of signaling factors by linkage to ubiquitin ligases, and broad inhibition of host gene expression through inhibition of mRNA export, splicing and/or translation.

Increasing human population density and travel enhance the probability of viral pandemics. In the case of SARS-CoV-2, there was a ∼30 day lag time between determination of the viral sequence and the full appreciation of its pandemic potential based on epidemiology. It would be ideal to rapidly estimate the pandemic potential of a virus based on its sequence. We envision a reductionist approach, in which gene activities are measured, mechanisms are inferred, and a calculation of pathogenicity is made. While this goal has not yet been realized, essentially all new viruses fall into known families, in which the pathogenicity of some members is known. By assaying a new virus in parallel to similar viruses of known pathogenicity, it should be possible to rapidly and accurately assess the danger of a new strain *in vitro*.

With this issue in mind, we generated a suite of ‘high-content’ cell-based assays which reveal the ability of viral proteins to inhibit the innate immune system. These assays can be employed within a month or less, enabling rapid identification of pandemic potential for newly detected viruses. Medium and high-throughput ‘high content’ cell-based screens have been successfully used to identify drugs that affect cellular pathways or phenotypes agnostic to specific protein targets. For example, Kau et al. (2003) executed a cell-based screen for modulators of the nuclear/cytoplasmic localization of a Forkhead transcription factor based on immunostaining; this effort ultimately led to the development of selinexor, an approved anti-cancer drug (Syed et al., 2019). Similarly, Mayer et al. (1999) screened compounds for the ability to arrest cells in mitosis using a cell-based assay for enhancement of mitotic phosphorylation sites; that effort led to development of Eg5 inhibitors, which entered clinical trials for cancer treatment (Rath et al., 2012).

To identify and understand the mechanisms that distinguish SARS-CoV-2 from other coronaviruses, we constructed mammalian and yeast expression vectors for genes encoded by SARS-CoV-2, SARS-CoV-1, MERS, 229E, NL63, HKU1, and OC43, and tested these genes for their ability to disrupt the innate immune system. Two assays are target-based screens that provide mechanistic insight into how individual viral genes disable the innate immune response measuring each gene’s effect on specific components of the Type I interferon response (IRF-3, NFkB, and STAT1 signaling pathways), and inflammatory gene expression (an NFkB-regulated reporter gene) (Koch et al., 2018). A third assay is a mechanism-agnostic screen that reveals the overall ability of viral proteins to successfully inhibit the immune response to a degree that supports viral replication (Brown et al., 2016). In doing so, we confirm the immune-suppressing role of certain coronavirus genes, implicate others, and report the particularly potent effect of SARS-CoV-2 on the innate immune system. In addition, we identify a novel protein encoded by SARS-CoV-2 which demonstrates potent immune-suppressing ability via the IRF-3 pathway.

## Results

We designed mammalian expression vectors for all proteins encoded by seven human-infecting coronaviruses: SARS-CoV-2, SARS-CoV-1, MERS-CoV, HCoV 229E, HCoV NL63, HCoV HKU1, and HCoV OC43 (Supplemental Information). The RNA of coronaviruses can be conceptually divided into two parts: (i) the 5’ two-thirds of the genome which encodes a ∼7,000 amino acid polyprotein termed ORF1ab that is cleaved into proteins NSP1-16, and (ii) the 3’ one-third of the genome which has multiple separate ORFs. This latter region encodes the universal proteins Spike, Envelope, Membrane, and Nucleocapsid, plus a number of small so-called “accessory” proteins that vary from virus to virus, some of which are encoded by overlapping ORFs. Of the three highly pathogenic coronaviruses, SARS-CoV-2 and SARS-CoV-1 are closely related, while MERS-CoV is somewhat related (Figure 1A). Four human-infecting coronaviruses (HCoV OC43, HCoV HKU1, HCoV NL63, and HCoV 229E) are quite distantly related; these viruses generally do not cause severe disease in healthy adults, although it is unclear whether their lack of lethality is because they do not suppress the innate immune system as severely as the SARS/MERS group or an alternative explanation such as suppression by pre-existing immunity to related viruses. The vectors were constructed by Twist Biosciences (Methods, Supplemental Information) and used directly for mammalian cell transfection, or individual viral genes were inserted into yeast for the yeast-mammalian cell fusion assay.

**Figure 1.**
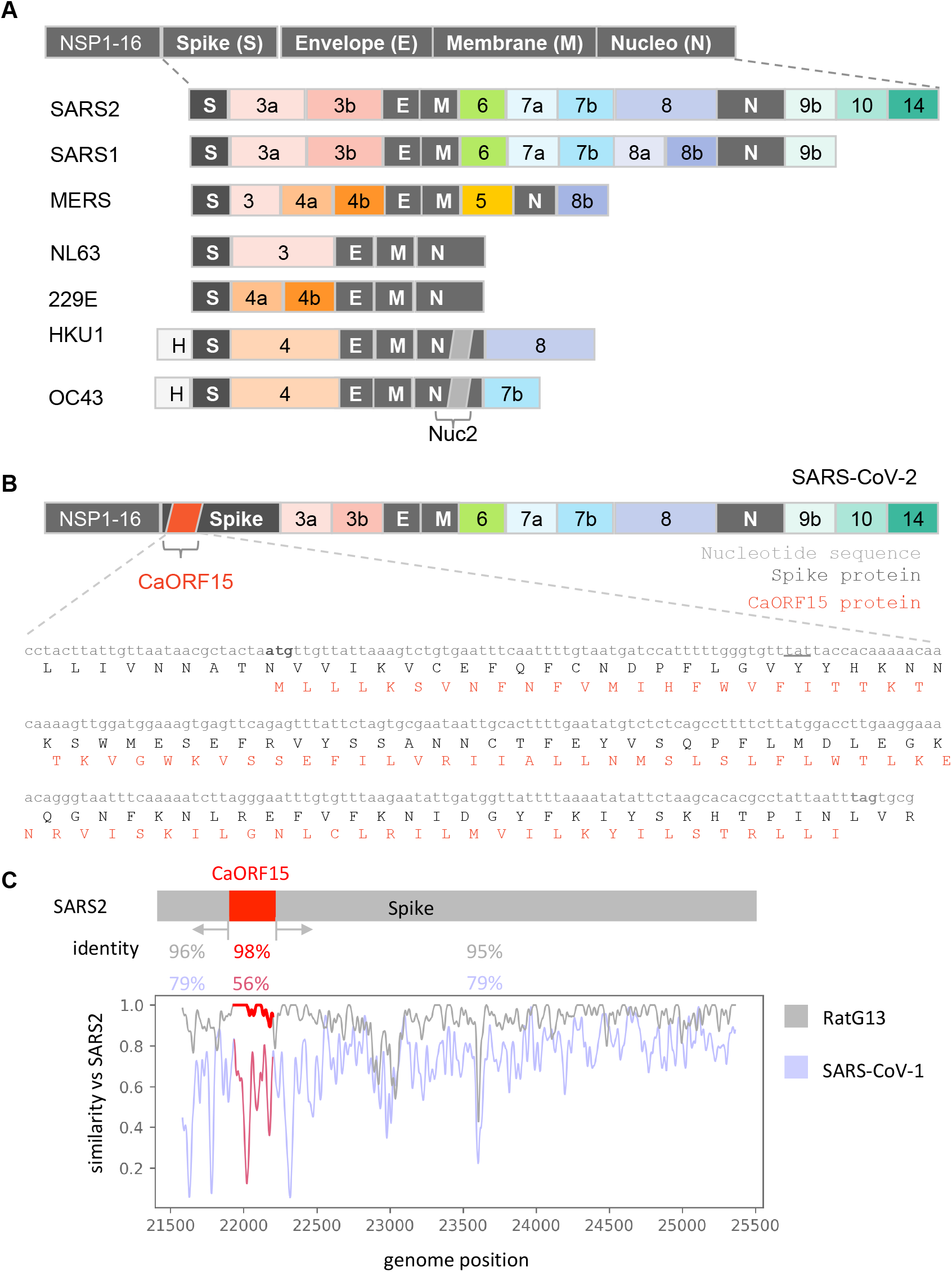
Identification of an additional ORF within the Spike coding sequence of SARS-CoV-2. **A.** Schematic of protein coding regions of the seven coronaviruses tested. The 5’ two-thirds of the genome encodes a polyprotein found in all coronaviruses which is cleaved into proteins NSP1-16. The 3’ one-third of the genome encodes the universal proteins Spike, Envelope, Membrane, and Nucleocapsid, plus several small proteins that vary from virus to virus. **B.** To identify candidate ORFs in SARS-CoV-2, we translated all six reading reading frames and identified a possible 87-amino acid ORF within Spike in an alternative reading frame. Scanning the sequences against known PFAM protein domains (ref PFAM database 2021 doi: 10.1093/nar/gkaa913) revealed a weak similarity to part of the “GTRA-like” PFAM domain (PF04138) (Guan et al., 1999; Gandini et al., 2017). **C.** Similarity plot for the SARS-CoV-2 genomic sequence within the Spike region as aligned with the bat coronavirus RaTG13 and SARS-CoV-1. The region encoding CaORF15 is more conserved than the other segments of the Spike coding sequence for RaTG13 and less conserved with respect to SARS-CoV-1.

To ensure we screened all SARS-CoV-2 genes, we scanned the genome (Wuhan-hu-1, MN908947) for potential ORFs by searching for similarity to known proteins. Enumerating all translations in the 6 reading frames and aligning to sequence profiles of protein families from PFAM (Mistry et al. 2020 doi: 10.1093/nar/gkaa913) (Methods), we identified a candidate ORF 15 (CaORF15) encoded in an alternative reading frame within the Spike sequence (Figure 1). CaORF15 is 87 amino acids long (by comparison, Orf9b is encoded in an alternate frame of Nucleoprotein and is 98 amino acids long). We predict CaORF15 to contain three transmembrane alpha helices 1-21, 36-55, and 64-87 (Hofmann & Stoffel 1993) and share similarity to the GtrA-like protein family (Guan et al., 1999; Gandini et al., 2017). The corresponding region in SARS-CoV-1 is very divergent (56% identity) and does not encode CaORF15. However, CaORF15 is encoded in the SARS-CoV-2-related bat virus RaTG13 and shares higher identity than the remainder of Spike (98% versus 93% respectively) (Figure 1C).

### Construction of cell-based assay for suppression of Type I interferon pathways

A striking feature of SARS-CoV-2 is the ability to infect humans and produce virus that can replicate and spread to other humans without inducing symptoms. Typically, ‘flu-like symptoms’ resulting from viral infection are a consequence of Type I interferon expression (Maher et al., 2007). Patients who receive interferon α for hepatitis or interferon β for multiple sclerosis experience flu-like symptoms from these proteins alone.

We therefore designed a cell-based assay to reveal the effect of individual coronavirus proteins on Type 1 interferon pathways. The innate immune response to viruses relies on cytokine signaling induced through the following cascade (Figures 2A and 2B): (i) out-of-place nucleic acids (e.g. dsDNA or dsRNA in the cytoplasm) are detected by pattern-recognition receptors (PRRs) (Hoffmann et al., 2015); (ii) PRRs initiate signaling pathways that cause translocation of the transcription factors Interferon Response Factor 3 (IRF-3) and Nuclear Factor κB (NFκB) to the nucleus; (iii) activated IRF-3 and NFκB in the nucleus induce the expression of Type I interferons and other cytokines to warn the surrounding cells of incoming viral attack, initiate the adaptive immune response, and attract specialized immune cells; and (iv) signaling by Type I interferon receptors on the surface of surrounding cells causes translocation of Signal Transducer And Activator Of Transcription (STAT) transcription factors to trigger a state of antiviral hypervigilance (Hoffmann et al., 2015).

**Figure 2.**
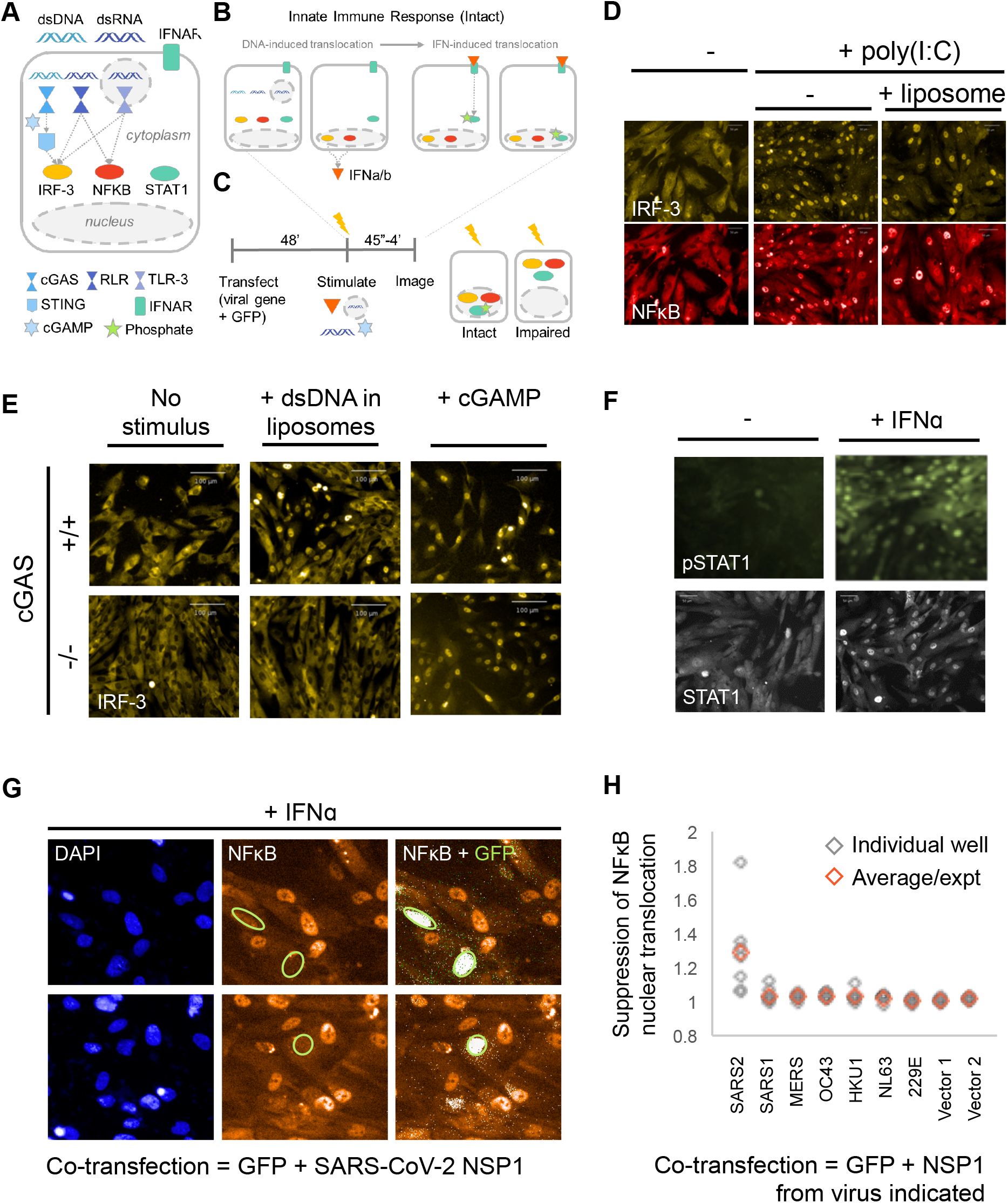
Construction of high-throughput antiviral innate immune response assay. **A.** Innate immune system signaling pathways that respond to viral infection. Viruses encode proteins that disrupt these pathways, by directly or indirectly inhibiting nuclear translocation or causing degradation of three key transcription factors: IRF-3, NFκB, and/or STAT1. **B.** Viral infection (i.e., presence of nucleic acids in an inappropriate cellular context) activates the transcription factors IRF-3 and NFκB. Together, IRF-3 and NFκB activate expression of type I interferons, which are secreted and act locally to stimulate the STAT1 pathway. IRF-3, NFκB and STAT1 all translocate into the nucleus during signaling. **C.** Experimental paradigm for testing virus genes for modulation of the IRF-3, NFκB, and STAT1 signaling pathways. BJ-5ta cGAS-/- cells are transiently transfected with a viral gene expression vector. The innate immune response pathways can be initiated *in vitro* by addition of cGAMP (a second messenger), extracellular double-stranded RNA (dsRNA), cytoplasmic dsRNA (by transfection), or interferon alpha (IFNα). Cells are then fixed and stained with antibodies against IRF-3, NFκB, STAT1, and/or pTyr701-STAT1, incubated with secondary antibodies, and then imaged. The effect of the viral protein on the innate immune response can be determined by quantifying the degree of nuclear accumulation. **D.** BJ-5ta cells treated with poly(I:C) or with liposomal poly(I:C) show translocation of both IRF-3 and NFκB into the nucleus. **E.** Knockout of the cGAS gene prevents transfection-mediated stimulation of IRF-3 translocation, while allowing downstream activation of IRF-3 via cGAMP. Double-stranded DNA in the cytoplasm activates the enzyme cGAS to create the cyclic dinucleotide cGAMP, which then acts on STING to activate IRF-3. Parental BJ-5ta cGAS+/+ cells show nuclear translocation of IRF-3 in response to transfected DNA and also to exogenous cGAMP (top), while BJ-5ta cells with a CRISPR knockout of cGAS show an IRF-3 response only after treatment with cGAMP (bottom). Thus, knockout of the cGAS gene allows assessment of elements of the STING/IRF-3 pathway without interference by transfected DNA. **F.** Treatment with IFNα causes STAT1 phosphorylation and translocation into the nucleus. **G.** Primary image data showing inhibition of NFkB translocation into the nucleus after polyI:C stimulation of BJ-5ta *cGAS^−/−^* cells co-transfected with SARS-CoV-2 NSP1 and GFP expression constructs. **H.** Results of image processing of sets of seven wells per condition, for cells transfected with NSP1 genes. See Supplemental Information for full details.

Combined activation of the transcription factors IRF-3 and NFkB is central to development of an antiviral response. Koch et al. (2018) recently developed conditions in which the nuclear/cytoplasmic localization of both proteins could be simultaneously visualized using non-cross-reacting primary and secondary antibodies. This system has been adapted for high-throughput screening to identify compounds that activated one or both factors (e.g., candidate anti-cancer or antiviral compounds), or inhibited the response to added cGAMP (candidate anti-inflammatory compounds). We adapted this screen for the present work.

Assay 1 was designed to test coronavirus genes for suppression of innate immunity via disruption of IRF-3, NFkB, and STAT1 (Figures 2 and 3). We used a non-transformed, tert-immortalized fibroblast cell line, BJ-5ta cells because they are thought to have an intact innate immune system. However, in normal cells, DNA transfection alone stimulates the cGAS/STING pathway (Figure 2E): cGAS binds to cytoplasmic DNA and produces the cyclic dinucleotide cGAMP as a second messenger that activates STING and downstream signaling steps (Xiao and Fitzgerald et al., 2013). We therefore generated a CRISPR knockout of cGAS (Methods). Because the coronaviruses are RNA viruses, removal of cGAS is not expected to be relevant in the present experiments; cGAS and cGAMP are inhibited or inactivated by certain genes of DNA viruses, such as poxin (Eaglesham et al., 2019), which cleaves cGAMP, but such an activity is not expected to benefit an RNA virus.

**Figure 3.**
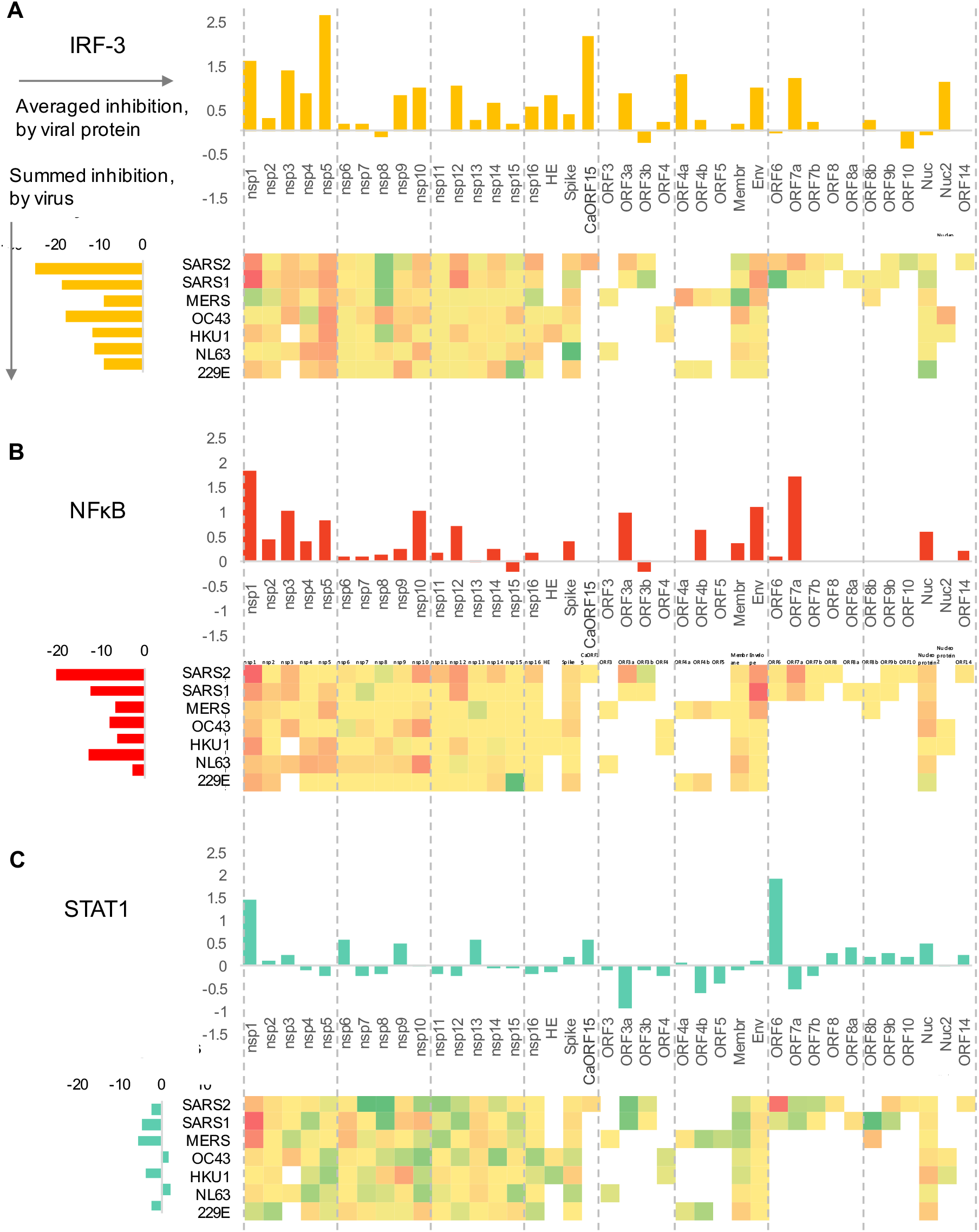
Inhibition of IRF-3, NFkB and STAT1 active nuclear levels by coronavirus genes. **A-C.** Statistically significant average inhibition scores from two to five experiments for each gene from each virus (e.g., Figure 2H) were combined. *Top:* Vertical bar graphs were generated by averaging the inhibition scores for a given gene across all viruses with that gene. *Left:* Horizontal bar graphs show the sum of all of the inhibition scores of genes from a given virus. *Bottom:* Heat map of average scores across all relevant stimuli in all experiments (each gene for each virus). Red indicates inhibition of innate immune signaling, green shows enhancement; white indicates a given gene is not present in a given virus; gray indicates a gene not tested. **A.** Inhibition of IRF-3 translocation and enhancement of degradation in cells stimulated by cGAMP plus low molecular weight polyI:C in lipofectamine plus free high molecular weight polyI:C. **B.** Inhibition of NFkB translocation and enhancement of degradation in cells stimulated by LMW-polyI:C in lipofectamine plus free HMW polyI:C. **C.** Inhibition of STAT1 translocation, enhancement of degradation, and formation of phospho-STAT1 in cells stimulated by interferon-alpha. See Supplementary Information for full details.

In the cGAS mutant cell line, DNA transfection-mediated stimulation of IRF-3 nuclear transport and Type I interferon were essentially undetectable (Figure 2E), and the functional transfection efficiency was increased (Supplemental Information). In one set of experiments, cells were simultaneously stained with antibodies to NFkB and IRF-3; in other experiments either anti-STAT1 or anti-phosphoSTAT1(Tyr701) were used (Methods). Treatment with either naked polyIC, lipofectamine/polyIC, or cGAMP was used to stimulate signaling via the TLR3, RIG-I/MAVS and STING pathways respectively; these stimuli cause a dramatic relocalization of IRF-3 and/or NFkB from the cytoplasm to the nucleus (Figures 2D and 2E). In other experiments, BJ-5ta *cGAS−/−* cells were treated with IFNalpha to initiate STAT1 phosphorylation and nuclear translocation (Figure 2F).

We therefore used the mutant cell line to conduct the following assay (Figure 2C): cells in 384-well plates were co-transfected with a viral gene expression vector and a GFP expression plasmid. Two days later, cells were treated with cGAMP, IFNα, or a form of polyI:C to stimulate an innate immune response. Based on the timing of the induced response (distinct for the various stimuli), cells were fixed, stained with antibodies directed against relevant transcription factors, and imaged. Fields of cells were scored by automated image processing for expression of the GFP marker, indicating functional transfection, and for the total amount and nuclear/cytoplasmic distribution of NFkB, IRF-3, STAT1, or pTyr701-STAT1. Evaluation of effects of virus protein expression is based on comparison of cell phenotypes of transfected versus untransfected cells in the same well and subsequent comparison to mock-transfected wells as well as virus gene-transfected wells that were mock-stimulated. Apparent ‘hits’ are identified and inhibition strengths are calculated (Figures 2G and 2H; Methods and Supplemental Information). Other virally encoded suppressors of these pathways generally work by either suppressing nuclear translocation of a given factor, or by inducing its degradation; both mechanisms would be detected by our image-processing calculations.

In the present analysis, the data were aggregated to reveal the most clear-cut observations (see Supplementary Information for details of the data processing and rationale for the methods used to aggregate the data). Enhancement of transcription factor degradation and blockage of nuclear import are both identified through these analyses. We summed data for different versions of a given gene from different viruses to draw increased confidence about inferred activities of those genes (vertical bar graphs in Figures 3A–3C).

### SARS CoV-2 genes most strongly modulate Type I interferon-related signaling pathways

Many individual coronavirus proteins inhibited the nuclear accumulation of IRF-3 (Figure 3A) and/or NFkB (Figure 3B). Proteins with the strongest effects were NSP1, NSP3, NSP5, and Orf7a; proteins NSP4, NSP10, NSP12, Env, and CaORF15 had more modest effects.

We determined which genes had consistent effects across coronaviruses. To make these inferences, data for each gene within each virus were first averaged across multiple experiments and summed across multiple input signal types to generate heat maps of all the results; these data were further aggregated by averaging the scores for individual genes across viruses (Figures 3A–3C; See Supplemental Information for breakdowns). This analysis revealed, for example, that NSP5 from all of the tested viruses had an inhibitory effect on IRF-3 signaling. The NSP10 and NSP12 genes from several different viruses also inhibited both IRF-3 and NFkB nuclear accumulation. While the effects of these genes are modest, the fact that we observe effects of these genes from multiple different viruses suggests that the effects are not due to noise in the assay.

In contrast to the broad suppression of IRF-3 and NFkB translocation, coronaviral proteins largely did not act upon the STAT1 pathway in our assay, with the noticeable exception of two proteins: NSP1 and ORF6 (Figure 3C). Both of these are known inhibitors of host functions: NSP1 binds to the ribosome and inhibits translation of a subset of host genes (Tidu et al., 2020), while Orf6 protein binds to and functionally inhibits Nup98, a nuclear transport protein that carries certain proteins into the nucleus (Miorin et al., 2020) and whose expression is induced by inflammatory stimuli (Enninga et al., 2002).

A number of viral proteins appeared to stimulate, rather than inhibit, the STAT1 pathway (Figure 3C). Such stimulation may represent situations where the innate immune system successfully recognizes expression of a foreign or deleterious protein, such as by poor protein folding that leads to creation of inflammasomes (Masters and O’Neill, 2011). Apparent increases in transcription factor levels, as detected by immunofluorescence, may also occur when a virus protein stabilizes the target host protein in an inactive state; for example, US3 of HSV-1 hyperphosphorylates IRF-3 and appears to stabilize this protein in the cytoplasm (Wang et al., 2013).

We next estimated the cumulative capacity of each virus to inhibit the innate immune system by summing the inhibitory effects of each individual gene within each virus (Figures 3A–3C; horizontal bar graphs). This analysis indicates that the genes from SARS-CoV-2, in aggregate, suppress IRF-3 and NFkB signaling more than the aggregate of genes from other viruses. Part of this effect is accounted for by the larger number of genes within SARS-CoV-2 (i.e., the less pathogenic strains have fewer C-terminal ORFs, see Figure 1A). However, the immune-suppressing proteins encoded by SARS-CoV-2 within this region also generally showed stronger effects than their SARS-CoV-1 homologues. In particular, Orf3a, Orf6, and Orf7a from SARS-CoV-2 showed stronger immune-suppressing effects (Figures 3A–3C). These proteins have no homologues in MERS-CoV, HCoV 229E, HCoV NL63, HCoV HKU1 and HCoV OC43, which encode a different set of accessory proteins that generally seem to be less active in our assays. Collectively, we conclude that the individual and cumulative ability of coronaviral proteins to suppress the Type 1 interferon-mediate innate immune response is strongest in SARS-CoV-2.

### NSP1 from SARS-CoV-2 strongly inhibits NFkB-driven gene expression

We determined which coronaviral proteins act by suppressing the expression of inflammatory genes within the host cell by a second cell-based assay to test whether coronavirus genes interfere with expression of a TNFα-inducible reporter element (Figure 4A). Specifically, we constructed a stable pool of HEK293 cells (which have a high transfection efficiency) with a DNA construct in which a degradation-tagged fluorescent mScarlet protein was expressed from an artificial promoter with five NFkB binding sites upstream of a ‘minimal CMV’ promoter element (Figures 4B and 4C; Supplemental Information). Cells were transiently co-transfected with virus gene and GFP expression plasmids, stimulated with TNF-α, and then assayed microscopically and by flow cytometry for expression of the mScarlet reporter in GFP-expressing and non-expressing cells (Figures 4B–4D).

**Figure 4.**
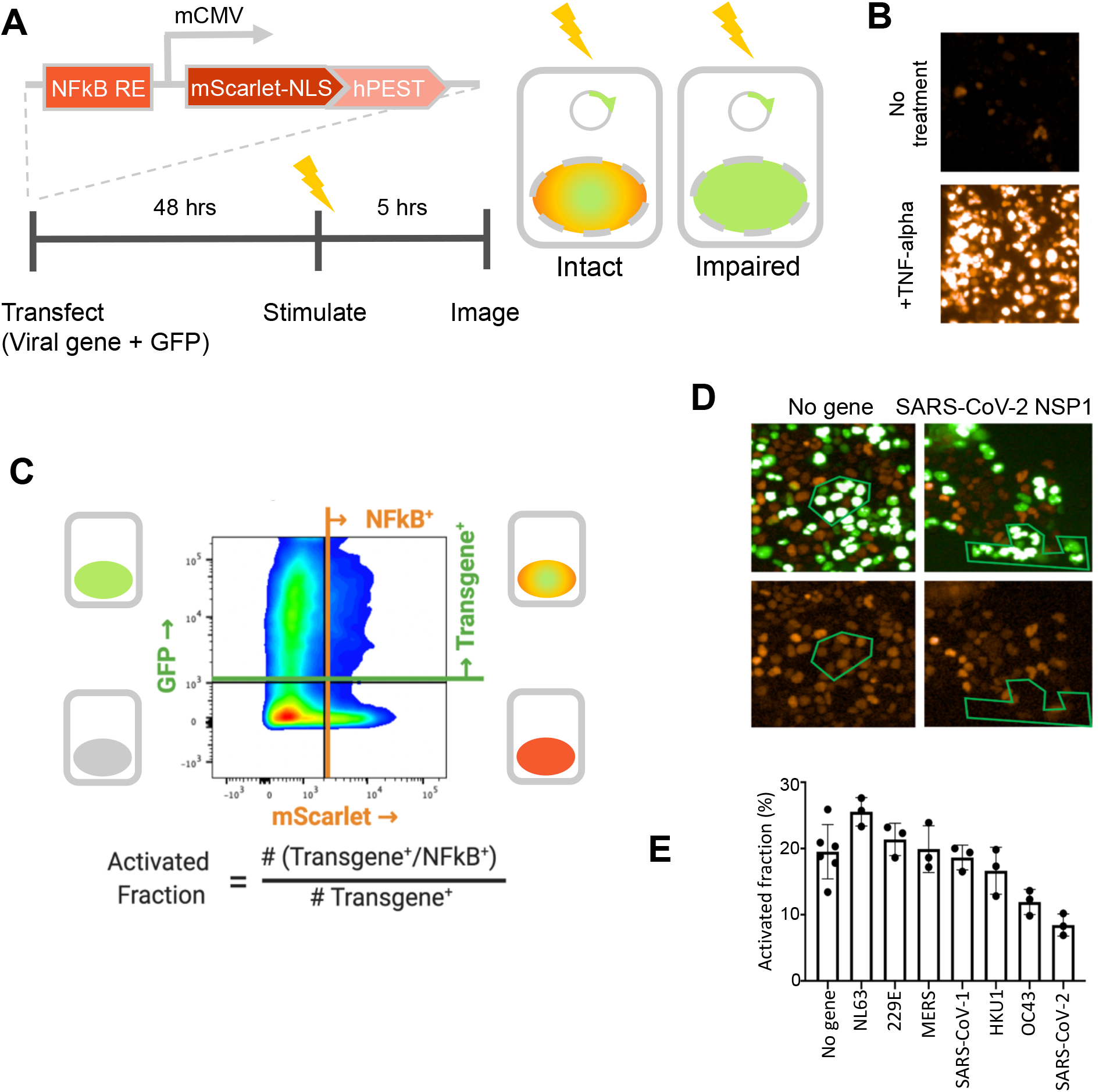
SARS-CoV-2 NSP1 strongly inhibits TNF-alpha activation of an NFkB reporter gene. A fluorescent NFkB reporter consisting of a 5x repeat of the NFkB consensus sequence (NFkB RE) upstream of a minimal CMV promoter (mCMV), and a human codon-optimized, nuclear-localized mScarlet (mScarlet-NLS) fused to an hPEST degradation tag (hPEST) was constructed and stably integrated into HEK293 cells. Reporter HEK293T cells were co-transfected with expression plasmids encoding GFP and a virus gene; 48 hours later, 5 ng/ml TNF-α is added; and 5 hours later, red and green fluorescence was quantified. **B.** HEK293T cells stably transduced with the reporter responded to human TNF-α by expression of mScarlet. **C.** Flow cytometry analysis of transfected, TNFα-treated cells. Upper quadrants are transfected cells. The upper right quadrant represents double-positive cells that are transfected and also express the inducible transgene. The activated fraction was calculated as the ratio of double positive cells to all GFP-positive cells. **D.** Reporter HEK293T cells were co-transfected with GFP and no-gene or SARS-CoV-2 NSP1 expression vectors, followed by TNF-α stimulation and imaging of red and green fluorescence. Green polygons highlight identical populations of cells in both images for representative comparisons. **E.** Activated fractions compared among NSP1s from several coronaviruses. Error bars represent the standard deviation of at least three technical replicates.

The NSP1 protein from several of the coronaviruses inhibited reporter gene expression, with the effect being strongest in SARS-CoV-2 (Figures 4D–4E). Figure 4D shows typical results for NSP1 of SARS-CoV-2: cells that express the GFP transfection reporter do not express the TNF-inducible mScarlet reporter, and vice versa. NSP1 is known to bind to the ribosome and disrupt translation of a subset of host mRNAs (Narayanan et al. 2015). Other genes showing inhibition of gene expression from more than one virus include NSP4, NSP6, NSP8, and NSP13, but the magnitude was much smaller than for the NSP1 genes (Supplemental Information). Notably, NSP1 was the only gene implicated in both the nuclear localization and gene expression assays. The difference in assay results may be because this transcription-based assay used HEK293T cells, which have a defective STING pathway and possibly other innate immune signaling defects (Sun et al., 2013).

### Cell-based assay for immune suppression-enabled viral replication

Because of the complex nature of human antiviral innate immunity, we developed multiple assays that could evaluate whether a viral protein was able to interfere with antiviral defenses. The assays above test the capacity of viral proteins to affect specific pathways involved in innate immunity. In another approach, we created a mechanism-agnostic test of innate immune suppression by viral genes, where the indicator of a viral protein that interferes with antiviral innate immunity is replication of a reporter virus. To do this, we built on our yeast-mammalian cell fusion system reported in Brown et al. (2016). There, we found that yeast spheroplasts could be fused to mammalian cells to deliver large DNAs to the mammalian nucleus, and also yeast-expressed proteins to the mammalian cytoplasm. In one variation of this experiment, a strain was constructed that contained the entire 155-kb herpes simplex type-1 (HSV-1) viral genome on a yeast centromeric (CEN) plasmid. When spheroplasted yeast were fused with HEK293 cells, viral particles were produced that could then productively infect other HEK293 cells. Typically, viral genomes in capsids and nucleocapsids have associated ‘non-structural’ proteins that are carried with the genome into the target cell; these proteins often suppress the innate immune system in the infected cell, facilitating replication without requiring viral gene expression in the cell. In the yeast-mammalian cell fusion experiment, such proteins are lacking. Fusion of such yeast with HeLa cells, which have a more intact innate immune system than HEK293 cells, did not allow viral replication. However, when yeast cells were further engineered to express the Ebola virus VP35 protein, fusion of such yeast with HeLa cells yielded infectious virus. As Ebola VP35 is a suppressor of intracellular innate immunity, this experimental paradigm establishes an assay for innate immune suppressors (Zampieri et al., 2007). We have used this assay system in the present work.

In the methodology used here, we first inserted most of the coronavirus genes described above into a galactose-inducible yeast expression vector in yeast by TAR cloning (Kouprina and Larionov, 2016) to construct one set of yeast for delivery of viral proteins. We constructed a second yeast strain in which the entire HSV-1 genome was inserted into a yeast *CEN* plasmid; this HSV-1 also contained a reporter gene in which green fluorescent protein (GFP) was expressed from a viral late promoter, so that viral replication could be easily assayed in a plate reader. The extent of viral replication as inferred from the GFP signal was highly correlated with direct titration of virus by a TCID_50_. Fusion of the HSV-1-containing yeast with HeLa cells does not generate live virus, but when the other yeast strain expresses Ebola virus VP35 protein, fusion of both yeast with HeLa cells yields replicating virus; this configuration serves as a positive control (Figure 5A; see Methods and Supplemental Information for details).

**Figure 5.**
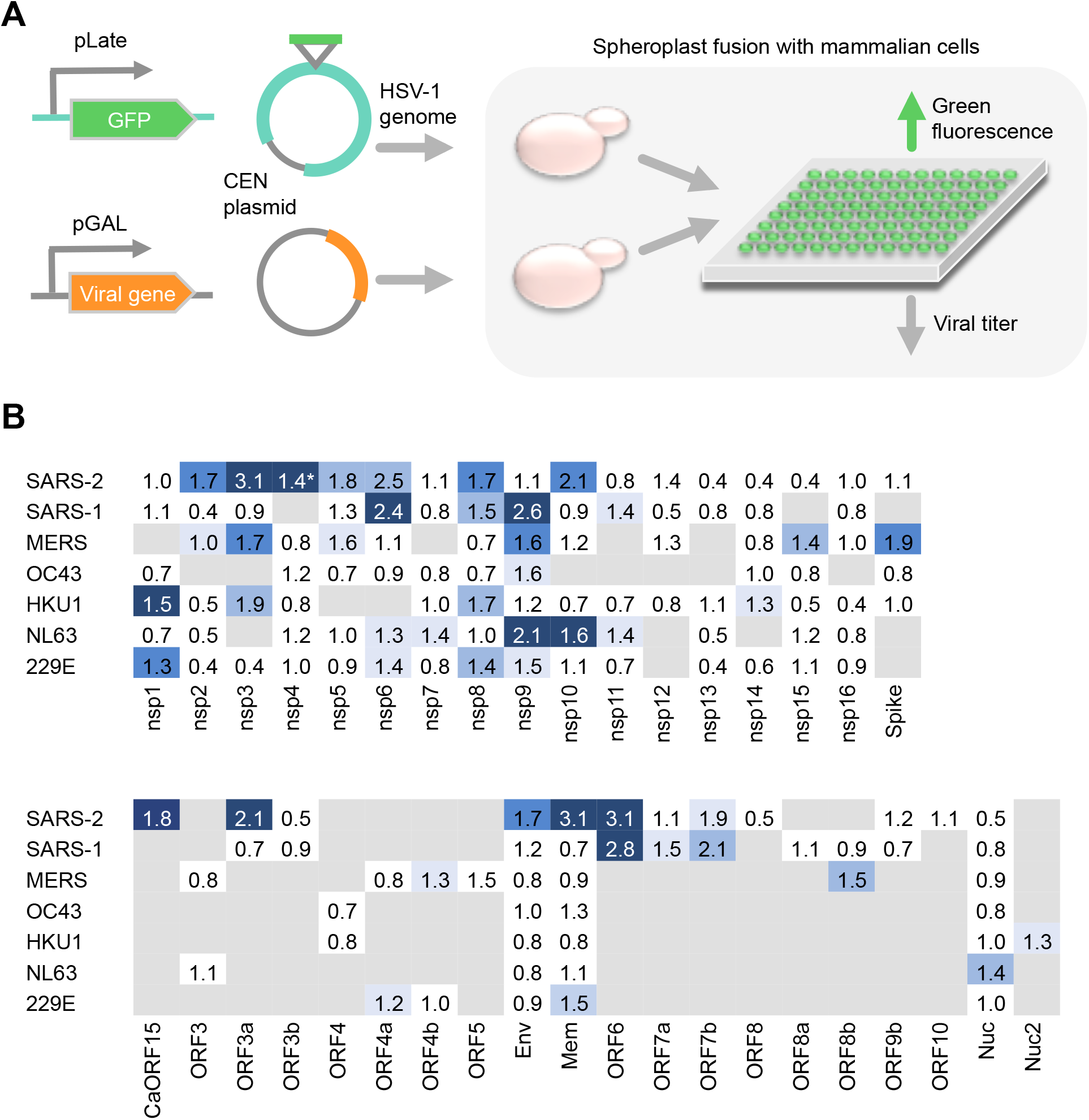
Assay for innate immune suppression based on protein delivery via yeast-mammalian cell fusion. **A.** Assay concept. A yeast cell with a plasmid expressing a virus gene from the *GAL1* promoter is induced with galactose to produce the viral protein in the yeast cytoplasm. A second yeast cell has a *CEN* plasmid containing the entire Herpes Simplex Virus-1 genome that also includes a GFP gene transcribed from a late promoter. The strains are mixed, spheroplasted, and fused with HeLa cells, which allows replication of the HSV-1 only if the viral protein delivered by the first yeast can suppress the innate immune system. The Ebola virus VP35 gene is a positive control (Brown et al., 2016). Immune suppression is measured via viral expression of GFP. **B.** Results from testing coronavirus genes. Numbers indicate the average fold increase in the GFP assay. Blue-highlights indicate a enhancement of viral replication (dark: p<0.05; medium: 0.05<p<0.1; light: 0.1<p<0.15; very light: 0.15<p<0.25); white indicates no stimulation detected; gray indicates that the genes were not tested or are not present. A number of genes score positively in this assay and also in the assays of Figure 2–3, including NSP3, NSP5, ORF6, and CaORF15.

The virus genes with the strongest effects were, in rough order, ORF6, ORF7b, CaORF15, NSP8, NSP9, NSP6, NSP3, ORF5, ORF3a, and Membrane (Figure 5B). A number of these showed effects in the transcription factor nuclear accumulation assays, such as ORF6, CaORF15, NSP3, and ORF3a. The fact that proteins such as ORF7b, NSP6, NSP8, NSP9, ORF5 and Membrane were clear positives in this mechanism agnostic assay but not in the mechanism-specific assays may point to the complex nature of both the innate immune system and the process of HSV-1 infection, which involves 77 viral proteins. Conversely there were proteins with no or minimal activity in the yeast fusion assays that were clearly positive in the other assays. NSP1 from SARS-CoV-2 and SARS-CoV-1 were negative in the yeast fusion assay, while the seemingly weaker NSP1 proteins (based on the transcription factor nuclear accumulation assays; Figure 4) from HKU1 and 229E were positive; this might be rationalized if the translation-inhibition function of the more potent versions of NSP1 interferes with expression of HSV-1 genes. The same reasoning could apply to other genes, for example proteins involved in coronavirus RNA processing and could interfere with HSV-1 gene expression. Thus, positive results obtained in the yeast-based assay may be more meaningful than negative results.

## Discussion

In this work, we characterized the ability of the proteins encoded by seven coronaviruses to suppress host intracellular innate immune signaling pathways using high-content, medium-throughput cell-based assays. The first assay examined the nuclear localization of the transcription factors IRF-3, NFkB, and STAT1, which play key roles in the elaboration of the Type I interferon response to viral infection, and which are major targets for inhibition by other viruses (Figure 2; Hoffmann et al., 2015). Viruses act on these pathways by a variety of mechanisms, such as specific inhibition of upstream signal transduction proteins, general inhibition of nuclear import, and enhancement of degradation. Our assays involved transfection of a non-transformed cell line with expression vectors encoding each gene followed by addition of a stimulator of innate immune signaling, and then immunofluorescence staining and automated image processing. We found that:

1. Localization of IRF-3 and NFkB to the nucleus was inhibited by a number of coronavirus genes, including known innate immune suppressors such as NSP5, NSP3, Orf7a and NSP1 (Freitas et al., 2020; Lei et al., 2018), and proteins not previously known to have such an activity, including CaORF15 and NSP9.
2. Two genes were found to have an impact on STAT1 activation: NSP1 and Orf6. Other groups previously found that that NSP1 binds to the ribosome and inhibits translation of a subset of host proteins (Tidu et al., 2021), and that Orf6 protein inhibits the nuclear protein import factor Nup98 (Frieman et al., 2007). Our results are consistent with these observations.
3. In the 3’ region of the coronavirus genome, which encodes accessory proteins that vary among the coronaviruses, SARS-CoV-2 proteins are more potent in their inhibition of innate immune signaling than the corresponding SARS-CoV-1 protein (Figure 3).
4. SARS-CoV-2 appears to encode a novel protein (“CaORF15”), which is predicted to have three transmembrane domains. CaORF15 is encoded in an alternate reading frame within the Spike coding sequence, and is not found in SARS-CoV-1 but is present in the more closely related bat virus RaTG13. CaORF15 inhibits IRF-3 nuclear accumulation particularly through the STING pathway (Figure 3, Supplemental Information) and also suppresses the innate immune system in the yeast-mammalian cell fusion assay (Figure 5).
5. When the inhibitory effects of coronavirus genes are examined in aggregate, SARS-CoV-2 appears to have more potential to suppress the innate immune system. This is based on summation of effects on IRF-3 and NFkB nuclear localization (Figure 3A and 3B), the strength of NSP1 inhibition of NFkB-mediated gene expression (Figure 4), and the genes showing effects in the yeast cell fusion assay (Figure 5).

NSP1 strongly blocks the activation of a TNF-inducible expression of a reporter construct with multiple NFkB binding sites (Figure 4). We found that only the NSP1 proteins from the various viruses strongly inhibited transcription of this reporter. NSP1 from SARS-CoV-2 had the strongest inhibitory effect in the TNF-inducible reporter assay (Figure 4F); this effect may contribute to SARS-CoV-2’s ability to blunt the host response. Similar transcription-based screens showed SARS-CoV-2 NSP1 inhibition of the IFN-beta promoter, which also depends on NFkB for activation (Lei et al 2020).

Our pathway-agnostic assay also identified NSP1, NSP3, CaORF15, and ORF6 as likely suppressors of innate immune signaling. Yeast containing both an attenuated HSV-1 genome and expressed coronavirus protein are fused with a mammalian cell; fusion of the yeasts with HeLa cells yields essentially no virus unless an active innate immune suppressor is also delivered, which then allows replication of the HSV-1 genome and production of live virus. The assay thus mimics an early step of virus infection, wherein ‘non-structural’ proteins are typically bound to the viral nucleic acid and are co-delivered. Such non-structural proteins often suppress the innate immune system; Ebola virus VP35 is an example (Kuhl and Pohlmann, 2012). This assay may detect proteins that act by distinct mechanisms from the pathway-focused tests. For example, NSP8 and NSP9, which were not detected in the other assays, score as positive in this test. Conversely, proteins that strongly inhibit host functions, such as SARS-CoV-2 NSP1, may interfere with HSV-1 replication in this assay and thus score as negative.

Viral proteases often act on virally encoded proteins and host proteins involved in innate immunity (Lei and Hilgenfeld, 2017). The NSP5 gene from each of the seven viruses tested had an inhibitory effect on IRF-3, and NSP5 from SARS-CoV-1, MERS, HKU1 and NL63 had an inhibitory effect on NFkB. In agreement with our findings, Freitas et al. (2020) predicted that the NSP5 papain-like protease would suppress innate immune signaling based on its cleavage and reversal of ubiquitin and ISG15 modifications. NSP3 is a large protein with many domains, including a papain-like protease. We also found that NSP3 genes from most of the viruses tested had inhibitory effects on IRF-3 and NFkB. NSP3 has a number of additional domains that may suppress innate immunity based on modulation of the ubiquitin ligase system (Lei et al., 2018).

We also found that NSP9, NSP10 and NSP12 had innate immune-suppressing activity. NSP9 from SARS-CoV-1, MERS, and NL63, promoted replication of HSV-1 in the yeast fusion assay, as did NSP10 from NL63 and SARS-CoV-2. NSP10 and NSP12 inhibited IRF-3 and NFkB. Lei et al. (2020) found that NSP12 inhibited the activation of a reporter gene containing the IFN-beta promoter. All of these proteins are involved with mRNA metabolism. NSP12 is the RNA-dependent RNA polymerase, so at a mechanistic level it is not apparent how this protein might modulate IRF-3 and NFkB. It is possible that, by analogy to influenza NS1 with its multiple surfaces that bind to unrelated host proteins, that NSP12 has surfaces that are not involved with RNA polymerization and can therefore carry out an unrelated function of titrating a host factor. Alternatively, these proteins may bind to host RNAs that express proteins involved in innate immunity.

As part of our preparation of expression vectors for coronavirus genes, we identified a new ORF in SARS-CoV-2, which we provisionally term “CaORF15” and which falls into the GTRA-like set of protein folds (PFAM family PF04138). The protein is encoded in an alternative open reading frame within the Spike coding sequence (Figure 1), specifically within the most N-terminal domain of Spike. Three lines of evidence suggest that CaORF15 encodes a functional protein and is not a fortuitous ORF. First, this region of the Spike coding region has an unusual sequence conservation pattern: both SARS-CoV-2 and the bat coronavirus RaTG13 encode this protein, and the coding region is more conserved at the nucleotide level than the rest of the aligned Spike sequence from these two viruses. In contrast, in the more distantly related viruses SARS-CoV-1 and M789 from pangolins that lack CaORF15, the nucleotide sequence is more divergent than for the rest of the Spike gene (Figure 1; Supplemental Information). These observations suggest a pressure to evolve CaORF15 and then to maintain it. Second, expression of this protein in BJ-5ta cells inhibited the nuclear translocation of IRF-3 in response to cGAMP (Figure 3, Supplemental Information). Third, delivery of this protein in the yeast fusion assay stimulates replication of yeast-delivered HSV-1. These results suggest that CaORF15 encodes a protein that functions to suppress the intracellular innate immune system. The presence of CaORF15 within the Spike coding sequences means that nucleic acid-based vaccines based on the natural Spike coding sequence – depending on their design -could also encode the CaORF15 protein, which may be immunosuppressive in a vaccine context. However, Pfizer’s vaccine is based on a codon-optimized version of Spike in which the AUG of CaORF15 is changed to ACG (WHO-INN, 2020).

In sum, we characterized the behavior of proteins from seven different coronaviruses in three different assays for suppression of intracellular innate immune signaling. We found that a number of proteins showed inhibitory activity. At least two of these, NSP9 and SARS-CoV-2 “CaORF15,” do not appear to have been previously identified as such.

Innate immune suppression may correlate with asymptomatic spread rather than pathogenicity per se. Infection with MERS, for example, is fatal much more often than infection with SARS-CoV-2 or SARS-CoV-1; here the severe symptoms, presumably a result of a robust Type I interferon response, prevent spread of the virus within the human population. In carrying out these experiments, we sought to address whether the pandemic potential of a virus could be estimated based on medium-throughput analysis of an entire viral genome for suppression of the human intracellular innate immune system.

Emerging viruses generally fall into known families. To set the stage for inferring pandemic potential in future emerging viruses, we compared the genes of SARS-CoV-2 with several other coronaviruses of known pathogenicity. The results of this analysis indicate that SARS-CoV-2 genes cumulatively appear to have a greater potential for immune pathway suppression than other coronaviruses, including SARS-CoV-1. This is admittedly an approximate statement, since the relative importance of each gene in these viruses is not generally known, but the data collectively are likely to suffice for a rapid assessment of whether to initiate a vaccine program. Taken together, these results suggest that rapid testing of viral genes in assays for innate immune suppression, performed with genes from related viruses, could be used for early-stage evaluation of the pandemic potential for emerging viruses.

## STAR METHODS

### RESOURCE AVAILABILITY

#### Lead Contact

Further information and requests for resources and reagents should be directed to and will be fulfilled by the Lead Contact Pamela Silver.

#### Materials Availability

Plasmids and strains are available upon reasonable request.

#### Data and Code Availability

All image datasets generated during this study are available upon request via OMERO and Dropbox.

### EXPERIMENTAL MODEL AND SUBJECT DETAILS

#### Searching for candidate ORFs

In early 2020, when annotation of the SARS-CoV-2 genome was poor, we chose to search the raw sequence (Wuhan-hu-1, MN908947) for any overlooked open reading frames. First, we translated in all six frames and collected all 399 translations longer than or equal to 10 amino acids (Supplementary Table 1). We reasoned that genuine proteins may show similarity to established protein families and therefore aligned all 399 translations against PFAM sequence profiles using HMMscan (Mistry et al. 2020 doi: 10.1093/nar/gkaa913; Eddy 1998 doi: 10.1093/bioinformatics/14.9.755).

Of the translations, 15 correspond to known SARS-CoV-2 proteins and each resulted in significant alignments to one or more PFAM profiles (all E-values < 1 E-14). At the time, ORF14 was missing from available SARS-CoV-2 annotation, nevertheless our approach identified the translation as a likely protein because of significant alignment to PF17635, now also called bCoV_Orf14 (E-value 6.2 E-35). In addition, while ORF3b is split in SARS-CoV-2 relative to SARS-CoV-1 by an early stop codon, we found alignments to the resulting fractional translations (E-values = 0.0024 and 0.17). There remained 11 candidate ORFs not known to be SARS-CoV-2 proteins and that aligned to some PFAM profile (E-values=6.4 E-5 to 0.94). Of these candidates, one stood out as the longest (87aa) and most significant (E-value = 6.4 E-5) by a large margin. We dubbed this translation “Candidate ORF 15” (CaORF15) and decided to test the sequence in our assays. CaORF15 encodes 87 amino acids at genome coordinates 21936-22199, within Spike protein coding region.

#### Conservation analysis of CaORF15

To find out whether CaORF15 is conserved in other coronaviruses, we performed a MAFFT alignment of the SARS-CoV-2, SARS, pangolin PCoV GX-P3B, and RatG13 genomes (MN908947, AY274119, MT072865, MN996532 respectively). We then computed the identity of the aligned CaORF15, and surrounding regions of Spike, versus nucleotides in SARS-CoV-2 (Fig 1C, top). We found that RatG13 encodes an amino acid sequence 93% identical to CaORF15 in SARS-CoV-2, whereas SARS-CoV and PCoV GX-P3B do not. Instead, SARS-CoV and PCoV GX-P3B encode shorter sequences (29aa and 51aa) with low identity to CaORF15 (24% and 25% respectively). For visualization, we scanned a 40-nt window over the aligned Spike regions, computing the non-gapped identity of each genome versus that of SARS-CoV-2 (Fig 1C, bottom). To emphasize local identity, we smoothed the windows by weighting matches with a centered normal distribution (sigma=8).

#### Preparation of mammalian expression plasmids containing virus genes

Genes encoding proteins from the seven coronaviruses SARS-CoV-2, SARS-CoV, MERS-CoV, HCoV 229E, HCoV NL63, HCoV HKU1, and HCoV OC43 were ordered from Twist Biosciences (Twist) or Integrated DNA Technologies (IDT). Sequences are listed in the Excel file [Table SX]. Coronavirus gene fragments corresponding to proteins NSP2-16 post-cleavage proteins were designed with added start codons for individual expression; normally these proteins are generated from cleavage of the NSP1-16 polyprotein (Fehr and Pearlman 2015). DNA sequences between 300 and 5000 bp were obtained from Twist Biosciences in a custom-onboarded vector based on pSecTag2 [Addgene #V90020], but lacking the signal sequence of that vector. Specifically, the methionine start codon deriving from pSecTag2 was also the start coding for the cloned viral genes. The same methods were used to construct expression vectors for controls: NS1 protein from influenza A virus [accession # NP_040984.1], the VP35 protein from Ebolavirus [accession # NP_690581.1], the V protein from parainfluenza virus 5 [accession # YP_138513.1], the V protein from measles virus [accession # YP_003873249], and the M protein from vesicular stomatitis virus [accession # NP_041714]. DNA sequences under 300bp were ordered from IDT with adapters for Gibson cloning using the same vector. The only genes over 5000 bp were the nsp3 gene fragments from SARS-CoV-2, SARS-CoV, MERS-CoV, HCoV HKU1, and HCoV OC43. These were ordered as clonal genes in two fragments, referred to as nsp3-1 and nsp3-2, and assembled with Gibson assembly into the wild-type full-length gene in pSecTag2. Most cloned genes were transformed into chemically competent DH5α *E. coli* (New England Biolabs C2987I). The NSP3 genes were difficult to construct as whole genes using DH5α, so we used *E. coli* PY1182 (an MM294-derived strain with a *pcnB80* mutation to reduce copy-number of ColE1 vectors; a gift of R. Losick) as a host.

DNA for transfection was prepared according to the QIAGEN endotoxin-free maxi-prep manufacturer protocol. After preparation, purified DNA was diluted in TE buffer to 200 ng/µL, aliquoted and stored at −20 °C.

#### High-Content Screening of IRF-3, NFkB and STAT1 signaling in BJ-5ta cells Culturing BJ-5ta human fibroblast cell line

BJ-5ta cells were purchased from ATCC (CRL-4001; https://www.atcc.org/products/all/CRL-4001.aspx) and cultured in the manufacturer recommended medium of 4:1 DMEM:M199 with 10% FBS (72% DMEM [ATCC 30-2002), 18% M199 [Thermo Fisher Scientific 11150059], 10% U.S Origin FBS (GenClone #25-514) with 10 µg/mL hygromycin B ([Invivogen ant-hg-1). Cells were cultured in T-25, T-75, and T-150 cell culture-treated flasks with vented caps (Corning) at 37 °C and 5% CO_2_. Cells were passaged every 3-4 days at 70-90% confluency to 30% confluency.

#### Seeding BJ-5ta cells into 384-well plates

BJ-5ta cells were lifted from T-150 culture flasks using 0.25% trypsin (VWR 45000-664) for 5-10 minutes, then the trypsin was quenched with 2x volume of culture medium and transferred to 50 mL Falcon tubes. The cell suspension was centrifuged in a swinging-bucket rotor at 300g for 6 minutes at room temperature. The supernatant was discarded by aspiration and cells were resuspended in a small amount of culture volume and counted with a Bio-Rad TC20 Automated Cell Counter. Cells were diluted to 60,000/mL and 40 µL of diluted cell suspension was added from a reservoir to all wells except the outer row (i.e. rows B-O, columns 2-23 were used) of tissue-culture treated black CellCarrier-384 Ultra Microplates (Perkin Elmer 6057302) using a 12-channel electronic multichannel 200 µL pipettor [Sartorius]. Plates were then centrifuged at 200g for 4 minutes at room temperature and incubated overnight. Typically, about 2-5,000 cells per well are seeded, and about 200-800 are transfected as defined by expression of GFP.

#### Co-transfection of virus gene plasmids and GFP into BJ-5ta cells

30-60 minutes prior to transfection BJ-5ta culture medium (4:1 DMEM:M199 with 10% FBS) was replaced with an equal volume of pre-warmed antibiotic-free transfection medium (4:1 DMEM:M199 with 20% FBS; 64% DMEM, 16% M199, 20% FBS). Transfection mixes were prepared according to manufacturer protocol (GeneXPlus, ATCC ACS-4004) with final concentrations of 1 µg DNA, 4 µL GeneXPlus in 100 µL of Opti-MEM I Reduced-Serum Medium. Briefly, GeneXPlus, plasmid DNA (200 ng/µL), and Opti-MEM I Reduced-Serum Medium (ThermoFisher #31985062) were warmed to room temperature and vortexed gently. Plasmid DNA was aliquoted into sterile microcentrifuge tubes at a ratio of 3:1 virus gene plasmid : GFP-containing plasmid. Opti-MEM was quickly mixed with GeneXPlus and the appropriate volume was added to each DNA aliquot and mixed briefly by gentle pipetting. GeneXPlus:DNA complexes were formed at room temperature for 15-20 minutes. Transfection mixtures were then added to each well at 10% final volume (4.4 µL transfection mixture was added to 40 µL transfection medium). Plates were centrifuged at 200 rcf, 4 minutes at room temperature to collect all transfection mixture into the medium and briefly mixed by tilting plate back and forth. Cells were incubated with transfection mixture at 37 °C, 5% CO_2_ for 24 hours to allow DNA to enter cells. Transfection medium was then exchanged for fresh culture medium and cells were further incubated for another 24 hours prior to stimulation with innate immune stimuli and fixation as described above.

#### Homozygous knockout of cyclic GMP-AMP synthetase (cGAS) in BJ-5ta cells

CRISPR was used to introduce a frameshift mutation at position 13 of exon 1 of cyclic GMP-AMP synthetase (cGAS) in BJ-5ta cells. Three different Synthego-designed guide RNAs were each co-transfected with Cas9-containing plasmid (Synthego) into BJ-5ta cells (ATCC CRL-4001) using Lipofectamine 3000. After 48 hours, samples were removed from each knockout pool for Inference of CRISPR Edits (ICE) analysis to assess gRNA efficiency, which was between 1% and 6%. The knockout pool with 6% gRNA efficiency (gRNA 2) was diluted to a density of 0.5 cells/100uL and plated into 96-well plates for clonal expansion. Colonies grown from a single cell were visually identifiable after 3 weeks. After 8 weeks, the cGAS locus was sequenced in each clonal colony to identify colonies with homozygous indels. One homogygous knockout colony was identified from 20 screened. Homozygous knockout in the successful colonies was confirmed via Western Blot for cGAS protein.

#### Stimulation of innate immune signaling

The BJ-5ta *cGAS^−/−^* cell line was used for these experiments. This cell line showed a very reduced level of background innate immune signaling that otherwise resulted from introduction of transfecting DNA. In addition, the transfection efficiency of these cells was improved relative to the parental BJ-5ta cells. Cells intended for stimulation with poly(I:C) HMW or liposome-encapsulated poly(I:C) LMW were primed 48 h in advance of stimulation by treating with interferon α1 (Cell Signaling #8927) or interferon α2b (PBL Assay Science #11100-1) at 50 ng/mL (final concentration in the well 5 ng/mL). 24 h after treatment with interferon, the cell medium was exchanged to remove external interferon from the cell environment. Different innate immune stimuli were applied to cell medium at 10% culture volume as follows. Low molecular weight (LMW) Poly(I:C) (Invivogen #tlrl-picw) at a concentration of 100 ng/µL (final concentration in the well 10 ng/µL) complexed with Lipofectamine 3000 (ThermoFisher #L3000015) in Opti-MEM reduced-serum medium (ThermoFisher #31985062) according to manufacturer’s instructions was used to stimulate RIG-I-Like Receptor (RLR) activity by incubation at 37 °C, 5% CO_2_ for 4 h. High molecular weight (HMW) Poly(I:C) (Invivogen #tlrl-pic) at a concentration of 1mg/mL (final concentration in the well 100 ug/mL) was used to stimulate TLR-3 activity by incubation at 37 °C, 5% CO_2_ for 2 h. 2’,3’-cyclic GMP-AMP (cGAMP) (Invivogen #tlrl-nacga23-5) at a concentration of 1 mg/mL (final concentration in the well 100 ug/mL) was used to stimulate STING pathway activity by incubation at 37 °C, 5% CO_2_ for 2 h. Interferon α1 (Cell Signaling #8927) or α2b (PBL Assay Science #11100-1) at a concentration of 50 ng/mL (final concentration in the well 5 ng/mL) was used to stimulate IFNAR activity by incubation at 37 °C, 5% CO_2_ for 45-50 min. Cell signaling was stopped by fixation as described below.

#### Cell fixation and immunofluorescent staining

Cells were fixed with 15 µL of 16% methanol-free formaldehyde (ThermoFisher #28908) added directly to the 45 µL of cell medium in the wells for a final fixation solution of 4% formaldehyde. After a 20 minute incubation, the 4% formaldehyde solution was aspirated and the cells were washed 3x with 60 µL PBS using an automated plate washer (BioTek EL406). Cells to be stained for phospho-STAT1 were further permeabilized with ice-cold 100% methanol (Sigma Aldrich #34860) and incubated at −20 °C for 10-15 minutes, then washed 3x with 60 µL PBS using an automated plate washer. All primary and secondary antibodies were diluted 1:400 in PBS containing either 2.25% bovine serum albumin [Millipore Sigma #A2058] or 5% normal goat serum (Abcam #ab7481) for blocking and 0.15% Triton X-100 (Sigma Aldritch #T8787) for permeabilization. Fixed cells were stained with 40 µL diluted primary antibody solution overnight at 4 °C. Cells were then washed 4x with 60 µL PBS using an automated plate washer and stained with 40 µL diluted secondary antibody solution with DAPI (ThermoFisher #D1306) added to a final concentration of 0.2 µg/mL. Cells were finally washed 4x with 60 µL PBS using an automated plate washer and sealed using impermeable black plate seals. If not imaged immediately, fixed and stained cells were stored at 4 °C for a maximum of 4-7 days.

Cells treated with interferon α were stained either for phospho-STAT1 (Cell Signaling Technology #9167) or for STAT1 (Cell Signaling Technology #14994). Cells treated with cGAMP, poly(I:C) HMW, or poly(I:C) LMW were stained simultaneously for IRF-3 (Cell Signaling Technology #11904) and NFκB (Santa Cruz Biotechnology #sc-8008). IRF-3, phospho-STAT1, and STAT1 primary antibodies were detected using an Alexa-Fluor 647-conjugated goat anti-rabbit IgG antibody (ThermoFisher #A21245). NFkB primary antibody was detected using an Alexa-Fluor 568-conjugated donkey anti-mouse IgG secondary antibody (ThermoFisher #A10037).

#### High-content imaging and image segmentation

Fluorescently stained plates were imaged on a PerkinElmer Operetta CLS High-Content Imaging System with a 20x, numerical aperture 0.75 objective. 20-25 sites were imaged in each well, covering 90-100% of the well. Each well was imaged for DAPI, GFP, and Alexa 647. Wells treated with cGAMP or poly(I:C) were also imaged for Alexa 568. Image segmentation was performed using Columbus software (PerkinElmer). Nucleus areas were identified with Columbus Method C based on the DAPI channel and cytoplasmic areas were assigned with Columbus Method D based on the Alexa 647 channel. Average intensity in the GFP, Alexa 647, and Alexa 568 (if applicable) channels was calculated for the cytosol and nuclear areas of each computationally identified cell. Single-cell results were exported from Columbus in CSV format and can be viewed at [insert Harvard Dataverse data ID here upon acceptance].

#### Data processing

Single-cell results were analyzed using a custom Python script which can be found at [insert GitHub URL here upon acceptance]. Briefly, nuclear objects identified by Columbus that correspond to cell debris and artifacts were eliminated based on nuclear morphology. For each transcription factor in each cell, Nuclear Localization (Nucleus intensity / Cytosol intensity) and Total Cell intensity (Nucleus intensity + Cytosol intensity) were calculated. Within each well, cells were sorted into GFP positive (GFP+) or GFP negative (GFP-) (as a proxy for expression of virus protein) based on average Nucleus GFP intensity. The GFP positive or negative cutoff was set at twice the median Nuclear GFP intensity (the median being within the distribution of the more numerous GFP-negative cells). Within each well, the average Nuclear Localization or Total Cell intensity was calculated for the GFP+ or GFP-subsets of cells. Subsequently, for each well, the average Nuclear Localization or Total Cell intensity for the GFP+ cells was normalized to the corresponding average for the GFP-cells to obtain a single normalized Mean Nuclear Localization or Mean Total Cell intensity.

Quality control was performed on a plate-by-plate basis as follows. If the mean of either of the two sets of controls containing no virus gene was outside the 20^th^-80^th^ percentiles of the plate data as a whole, the data for the aberrant control was discarded. If both no-gene controls were non-aberrant, the two sets of no-gene control data were combined for the following primary “fold change” calculation and normalization purposes. Additionally, for each individual set of 7 technical replicates, if any data point was more than 3 times the interquartile range higher than the 75^th^ percentile or lower than the 25^th^ percentile, it was removed from the analysis.

Within each plate, the mean and standard deviation of the 7 technical replicates of each virus gene/innate immune stimulus (one type of innate immune stimulus or mock stimulus per plate) combination were calculated. The fold change and corresponding significance of each set of 7 wells for a given virus gene that were treated with innate immune stimulus were calculated according to one of two different methods. Attached spreadsheet(s) in the folders “coronavirus_IRNF-raw” and “coronavirus_STAT-raw” contain these data of the 7 techical replicates. To further process the data, by a first method the fold change of virus gene-transfected wells treated with an innate immune stimulus was normalized relative to the fold change calculated for wells transfected with the identical vector lacking a virus gene. For the second method, virus gene-transfected wells were normalized by dividing by the mean of the mock-transfected wells for both the stimulated and the mock stimulated conditions. The fold change of normalized virus gene-transfected, innate immune-stimulated wells was then calculated relative to normalized virus gene-transfected, mock-stimulated wells. The more inhibitory of the metrics calculated by the two different methods was taken as the inhibitory score for that virus gene for that stimulus in that experiment. In addition, for subsequent calculations, fold change scores with p>0.1 were considered not significant and were represented as 0 for averaging and summing purposes in subsequent calculations.

Data were further aggregated as follows to generate data representations in Figure 3 and corresponding Supplementary Figures. For each gene from each virus, data from two to five experiments were averaged; in general, genes that showed no effects in the first two experiments were not re-tested. Inhibition scores were calculated from average fold-change scores by: (a) inverting the positive/negative sign, and (b) scaling the score to be in a range of roughly 0 to 1. The purpose of these calculations is simply to make the inhibition scores more intuitive. The rationale for changing the sign is that, for example, the response to an immune stimulus will be reduced relative to negative controls if a virus gene inhibits that response, so the sign of the fold change score will be negative. The rationale for re-scaling the scores is that the primary fold change scores are very small; this is in part a result of the calculations performed by the proprietary Columbus software, and likely in part results from the fact that we do not perform a background subtraction when assessing signal levels in the initial image processing. Thus, the primary fold change scores are thus highly artificial numbers; our confidence in their meaning results from the fact that differences from controls are statistically significant, that positive control genes such as parainfluenza virus V protein score correctly in our assays, and that coronavirus genes identified by others as major inhibitors of innate immune signaling also behave as expected in our assays.

Data represented by heat-maps in Supplementary Figures show the results for all of the coronavirus genes tested for each immunostimulatory input (cGAMP, high-molecular weight polyI:C, low molecular-weight polyI:C in lipofectamine, and interferon-alpha) and transcription factor output (IRF-3, NFkB, STAT1, and phospho-STAT1).

Data in Figure 3A-C were generated by summing these data across inputs: specifically, scores for IRF-3 were generated by summing the inhibition scores seen with cGAMP, hi-MW polyI:C, and low-MW polyI:C; and scores for NFkB were summed from hi-MW polyI:C, and low-MW polyI:C scores. These choices are based on the current understanding of innate immune signaling pathways: that cGAMP activates only IRF-3 while the forms of polyI:C stimulate both IRF-3 and NFkB. We considered the summation of these results to be legitimate because the outputs and corresponding staining antibodies are the same across values being summed. Data in Figure 3C result from summing results that derive from staining with an anti-STAT1 antibody and an anti-phospho-STAT1 antibody; these are different antibodies and could have different signal/noise properties and thus might yield quantitatively non-comparable results. However, we note that: (a) for each set of staining results, the inhibition scores fall into comparable ranges and that the scores for these two metrics are generally correlated; and (b) both ranges of inhibition scores can be bracketed by the empty vector negative control and the PIV5 V protein positive control.

#### Blinded testing and analysis

For each protein, at least three different samples of the corresponding prepped DNA were given to a third party, who randomized and blinded them. The samples were then tested, analyzed, and the identity of each protein was assigned based on comparison to unblinded samples run concomitantly.

#### High-Content Screening of TNFα signaling in HEK cells

##### Cell culture (Figure 4 and associated text in Results)

HEK293 [ATCC CRL-1573]and HEK293T/17 [ATCC CRL-11268] cells were maintained in DMEM + 10% fetal bovine serum at 37°C in humidified 5% CO_2_.

#### NFKB Reporter Lentivirus Construction

The fluorescent NFkB reporter construct was made by a Gibson assembly insertion of the mScarlet-hPEST sequence (fpbase.org) into the pGreenfire1 NFkB lentivector (System Biosciences). VSV-G pseudotyped lentiviral particles were made from the resulting lentivector. Briefly, 2*10^6^ HEK293T/17 cells (ATCC) were cotransfected with the NFkB reporter lentivector (2 μg) with pMD2.G (0.5 μg) and psPAX2 (1.5 μg) (Addgene), using Lipofectamine 3000 (ThermoFisher) according to the manufacturer’s instructions. After an overnight transfection, the media was replaced and cells incubated for 2 days. Lentiviruses were harvested and concentrated from 10 mL cell culture media to 1 mL concentrated lentivirus using Lenti-X concentrator according to the manufacturer’s instructions (Takara). Lentivirus aliquots were stored at −80C for further use.

The sequence of the resulting construct, “GF_NFKb-mSc-mPGK-Puro,” is in Supplemental Information.

#### NFKB Reporter Cell Line Generation

1*10^6^ HEK293 cells, seeded in a 6-well plate, were transduced with 200 μL of reporter lentivirus in the presence of 5 μg/mL Polybrene (Millipore) for two days. The NFkB reporter cells were then cultured in fresh media with 5 μg/mL puromycin (Gibco) for two days to generate a stable pool of reporter cells.

#### High-throughput NFkB Reporter Cell Screening

Stably-transduced HEK293 cells (passages 2-8 after transduction) were seeded in 100μL in 96 well plates at a density of 3*10^5^ cells/mL. The following day, co-transfection was performed with 75 ng virus gene and 25 ng GFP-NLS per well using Lipofectamine 3000 (Thermo Fisher Scientific) according to the manufacturer’s instructions. Each gene was transfected into three replicate wells. After 24 hours of transfection, the media + lipofectamine complexes was replaced with fresh media, and the cells were incubated overnight. The next day, cells were stimulated with 5 ng/mL human TNF-α (Peprotech) for 5 hours. The negative transfection control vector consisted of the empty backbone vector (No gene, or ‘EV’ for empty vector), while positive control vector consisted of a dominant-negative (DN) IkBα (S32A, S36A), a well-established inhibitor of NFkB.

#### Flow Cytometry

Following cytokine stimulation, the media was removed and the cells were stained with Zombie Violet Fixable Viability dye (Biolegend) to quantify live cells according to the manufacturer’s instructions. After removing the dye, cells were briefly trypsinized, collected by centrifugation, and resuspended in ice-cold FACS buffer (10% FBS in Dulbecco’s phosphate-buffered saline (DPBS) with 5 mM EDTA). Transfected cells were gated based on GFP fluorescence, and statistics on 10,000 transfected cells were collected per well for analysis.

#### Analysis

Flow cytometry data was analyzed in FlowJo. Data was plotted in Graphpad Prism. Each gene was assayed in 3 technical replicates and screened in at least two independent experiments.

#### HSV-1 Assay Protocol (Figure 5 and associated text in Results)

##### Preparation of yeast expression strains containing virus genes

We constructed yeast strains that express viral genes from the galactose-inducible *GAL1* promoter by using TAR cloning to combine the virus genes described above with a plasmid, pLDJIF15, that supplies the yeast promoter, 5’end material, 3’ end material, and a selectable marker. In this process, the virus gene is amplified from the mammalian expression vector, mixed with pLDJIF15, and the mixture transformed into an auxotrophic yeast. Prototrophic yeast strains will be those in which both DNA fragments have entered the cell and undergone homologous recombination to construct a circular, non-integrated plasmid. Thus, there is no intermediate E. coli carrier of the constructed plasmids.

DNA encoding virus proteins was amplified from the mammalian plasmids with primers D298 5’-aatatacctctatactttaacgtcaaggagCTATAGGGAGACCCAAGCTGGCTAGCCACC-3’ and D299 5’-aataaaaatcataaatcataagaaattcgcAGAAGGCACAGTCGAGGCTGATCAGCGGGT-3’ (lower case: sequences in the yeast expression vector pLDJIF15; upper case: sequences flanking the virus genes in the mammalian expression plasmids) using PrimeSTAR® Max DNA Polymerase (Takara R045A) in a 10 µl reaction. The common Kozak sequence used in the mammalian expression vectors was designed to be consistent with the S. cerevisiae consensus Kozak sequence, since this element is transferred into the yeast expression vectors. Mammalian expression plasmids (0.5 µl) described above were used as template for each reaction.

The backbone was amplified from pLDJIF15 with primers D300 5’-GGTGGCTAGCCAGCTTGGGTCTCCCTATAGctccttgacgttaaagtatagaggtatatt-3’ and D301 5’-ACCCGCTGATCAGCCTCGACTGTGCCTTCTgcgaatttcttatgatttatgatttttatt-3’ using PrimeSTAR® Max DNA Polymerase (Takara R045A). The amplified backbone was treated with DpnI (NEB R0176S) overnight at 37 °C and purified with NucleoSpin® Gel and PCR Clean-up Kit (MN 740609.5). One hundred ng of backbone and 1 µl of virus gene insert linear fragments from the 10 µl reaction were mixed and transformed into yeast (W303α: MATα ade2-1 ura3-1 his3-11 trp1-1 leu2-3 leu2-112 can1-00) via electroporation. Transformed yeast cells were plated on -TRP (Teknova C71731) plates for selection. Trp+ colonies were verified by colony PCR with primers: D66 5’-CAACCATAGGATGATAATGCGATTAG-3’ and D67 5’-TGAGAAAGCAACCTGACCTACAG-3’ using QIAGEN Multiplex PCR Kit (Qiagen 206143). At least 2 size-confirmed transformants were chosen and used in a spheroplast fusion experiment.

The sequence of pLDJIF15 is attached as a separate file.

For the HSV-1 assay-based experiments, we generally followed the protocol developed by Brown et al (2016) with the following variations. First, we used two yeast strains to fuse with the mammalian cells – one to deliver the viral protein

The process generally involves four steps, (1) preparing yeast spheroplasts, (2) preparing the mammalian cells, (3) fusing cells, and (4) analyzing the results. First HSV-1 genome containing yeast spheroplasts and the yeast spheroplasts containing the expression plasmid for each gene were prepared, 8 fusions were performed for each gene. To properly synchronize preparing yeast spheroplasts and the HeLa cells, yeast spheroplasts were prepared first and then frozen then thawed and fused with HeLa cells once ready. To prepare yeast spheroplasts; first, inoculate 20 ml of –Trp media with selected yeast strain in a 50 ml falcon tube for the viral test genes or -URA media for HSV-1 strain. Twenty-two different samples were prepared at once at 30°C and grown overnight. Yeast cells were resuspended in 40 ml YPG media (or YPD media for HSV-1 samples) by adding directly to tube. Then grown for 5-6 h at 30°C. Yeast cells were collected by centrifugation at 3,000 rpm for 3 min (50 ml Corning Inc. centrifuge tube) and re-suspend in 20 ml 1M Sorbitol. Cells were kept at 4°C 18h. Yeast cells were collected by centrifugation 3,000 rpm/3 min, and re-suspend in 20 ml SPEM. A 1:10 dilution of OD should be ∼ 0.9. 20µl ß-Mercaptoethanol (Sigma) and 20µl Zymolyase-20T [Zymolyase stock: 200 mg Zymolyase, 9 ml H2O, 1ml 1M Tris pH7.5, 10 ml 50% glycerol. Store at –20°C] were added. Yeast cells were incubated at 37°C for 23-25 min with gentle agitation. Checking the OD600 of the yeast suspension: 0.1ml+0.9 ml 1M Sorbitol; 0.1 ml+0.9 ml 2% SDS. Compare the OD readings of A/B ratio. Sheroplasting was stopped when the difference is 3-4 fold by adding 30 ml 1M sorbitol and gently mix (inverting). Cells were collected by centrifugation at 1800 rpm, 8 min, 4°C. Yeast spheroplasts were resuspended with 2 ml 1M Sorbitol. Yeast spheroplasts counted using the OD calculation worksheet to determine the amount of yeast cells needed for each fusion. Two samples worth of spheroplast solution calculated from worksheet was added to a 1.5 ml tube (typically, 40 – 400 µl for each 2-sample tube). Yeast spheroplasts resuspended with 200µl 1M Sorbitol +15% DMSO and stored at - 80°C to be used when the mammalian cells are ready.

To prepare Mammalian cells, HeLa cells were grown to 70 to 80% confluence, in T150 flasks with 35 ml DMEM media + 10% FBS + penicillin streptomycin and amphotericin B. Next 200 µl Colcemid was added to each flask and incubated overnight at 37°C. Flask was striked 3 times to dislodge loose cells. Media removed from HeLa cells and placed in an empty flask and cells trypsinized to remove cells from flask. Once cells have detached, cells were resuspended in saved spent media. To recover the frozen yeast spheroplasts: they were thawed on ice 5 min, 22µl 10X TC added to each tube and mixed, centrifuged yeast spheroplasts 4000 rpm 1 min at 4°C, resuspended yeast spheroplasts in 400 µl STC to yield 2 samples worth per tube and kept at 4°C until ready.

Next the cell fusion was performed. PEG solution (50% PEG 2000 + 10%DMSO in 75mM HEPES buffer) was prepared: 12.5g of PEG to a 50 ml tube, add 75mM HEPES to ∼22.5 ml mark, invert on a shaker for ∼30 min until dissolved, top with 75mM HEPES to 25ml, add 2.77 ml DMSO and mix. Mammalian cells counted using hemocytometer and add 3 x 10^5^ HeLa cells added to tube. HeLa cells were centrifuged down and resuspended in 500µl saved spent media. Next, 200 µl of yeast spheroplasts from previous step was added to tube, and cell mixture incubated 5 min at room temperature. Pelleted cells at 5000 rpm for 30 sec media removed. Next, 500µl of PEG solution was added and incubated for 5 min at room temperature. Fusion was ended by adding 1ml serum free DMEM and spinning down at 4000 rpm in a tabletop centrifuge for 30 sec, media removed, and 1ml of DMEM media supplemented with 10% FBS, penicillin streptomycin and amphotericin B added to the sample. Next, 500 µl of DMEM media supplemented with 10% FBS, penicillin streptomycin and amphotericin B was added to a 24 well dish, and 500 µl of the fused, recovered sample was added to the 24 well dish, after 4 h after cells had settled and reattached to plate so media was removed and replaced with 500 µl fresh media to remove excess yeast. Evidence of fused cells can be seen 96h post fusion for HSV-1 replication and Tecan using fluorescence protocol analyzed sample plate.

## ACKNOWLEDGMENTS

This work was supported by IARPA-FunGCAT **Cooperative Agreement W911NF-17-2-0092**. D. Burrill assisted with proofreading and organizing the manuscript.

## AUTHOR CONTRIBUTIONS

Erika J. Olson performed and supervised data collection for the screens in Figures 2 and 3, consolidated the data from these screens, performed data analysis and curation, and contributed to the writing; Tai L. Ng performed some the screens in Figures 2 and 3; H. Sloane Weiss constructed plasmids used in the experiments and performed some the screens in Figures 2 and 3; Timothy Z. Chang performed the screens in Figure 4; David M. Brown and Lin Ding performed screens and constructed plasmids for data in Figure 5; Yukiye Koide constructed the BJ-5ta cGAS knockout strain and expression vector design; Peter Koch, Yukiye Koide, Pia Mach, Tobias Meisinger, and Timothy Mitchison contributed to the screening methodology; Nathan Rollins, Joshua Rollins and Debora Marks performed to the sequence analysis and identification of CaORF15, Trenton Bricken and Colin Molloy contributed to the image processing; Yun Zhang contributed to the statistical analysis; Bridget N. Queenan contributed to the writing and figure design; Jeffrey C. Way was primarily responsible for funding acquisition and initial project conceptualization with assistance from Pamela A. Silver, John I. Glass, Timothy Mitchison, and Debora Marks; Pamela Silver was responsible for the overall project administration.

## DECLARATION OF INTERESTS

The authors declare that they have no conflicts of interest.

## SUPPLEMENTAL INFORMATION

Supplementary Information for Olson et al., “High-content screening of coronavirus genes for innate immune suppression reveals enhanced potency of SARS-CoV-2 proteins.”

### BLAST analysis of segments of the Spike coding sequence

To characterize the rate of evolution of the segment of Spike gene that also encodes CaORF15, we performed a BLAST alignment of segments of the SARS-CoV-2 Spike coding sequence with the Spike coding sequence from the coronaviruses RaTG13, pangolin M789, and SARS-CoV-1. Numerical values presented here correspond to the bar graph in Figure 1.

**Supplementary Figure 1.**
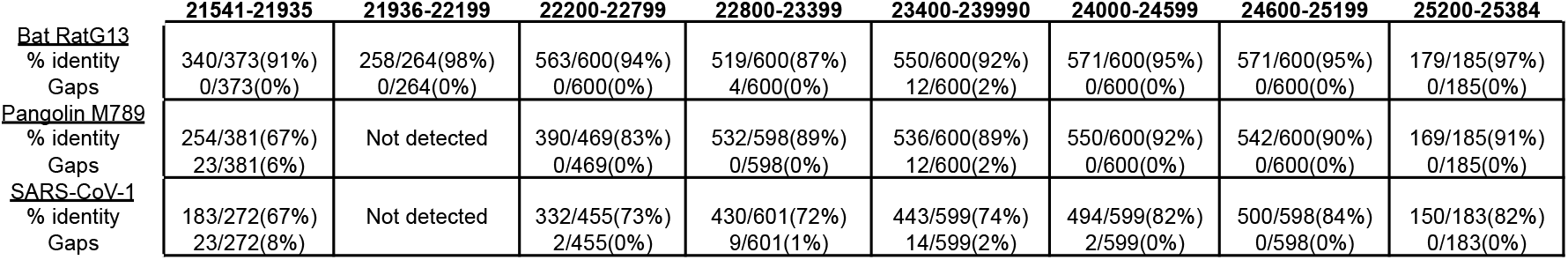

These results are comparable to Figure 1b of Boni et al. (2020), which indicate that the nucleotide sequence of the 5’-most region of the Spike gene in SARS-CoV-2 (which also encodes CaORF-15) is the most divergent region of the genome relative to the viruses SARS-CoV Guangzhou 2002 (HSZ-Cc), Zhejiang 2012 (Longquan_140),Hong Kong 2005 (HKU3-1), Zhejiang 2017 (CoVZC45), Zhejiang 2015 (CoVZXC21), Pangolin Guangxi 2017 (P1E), and Pangolin Guangdong 2019. However, this region is quite similar to Yunnan 2013 (RaTG13), which also encodes a CaORF15.

### Conservation of CaORF15 in SARS-CoV-2 strains

We then asked whether caORF15 was conserved among current variants of SARS-CoV-2 by examining the multiple sequence alignment of SARS-CoV-2 genomes provided by GISAID (Elbe & Buckland-Merret 2017 doi: 10.1002/gch2.1018). As of January 2021, of 276,082 human SARS-CoV-2 samples, 98% contain intact caORF15 and 92% have wholly identical nucleotide sequence to caORF15 to Wuhan-hu-1.

### Breakdown of effects of coronavirus genes on innate immune signaling

The following Figures are constructed in the same manner as Figure 3A-C, but represent the data for individual inputs and outputs.

**Supplementary Figure 2A.**
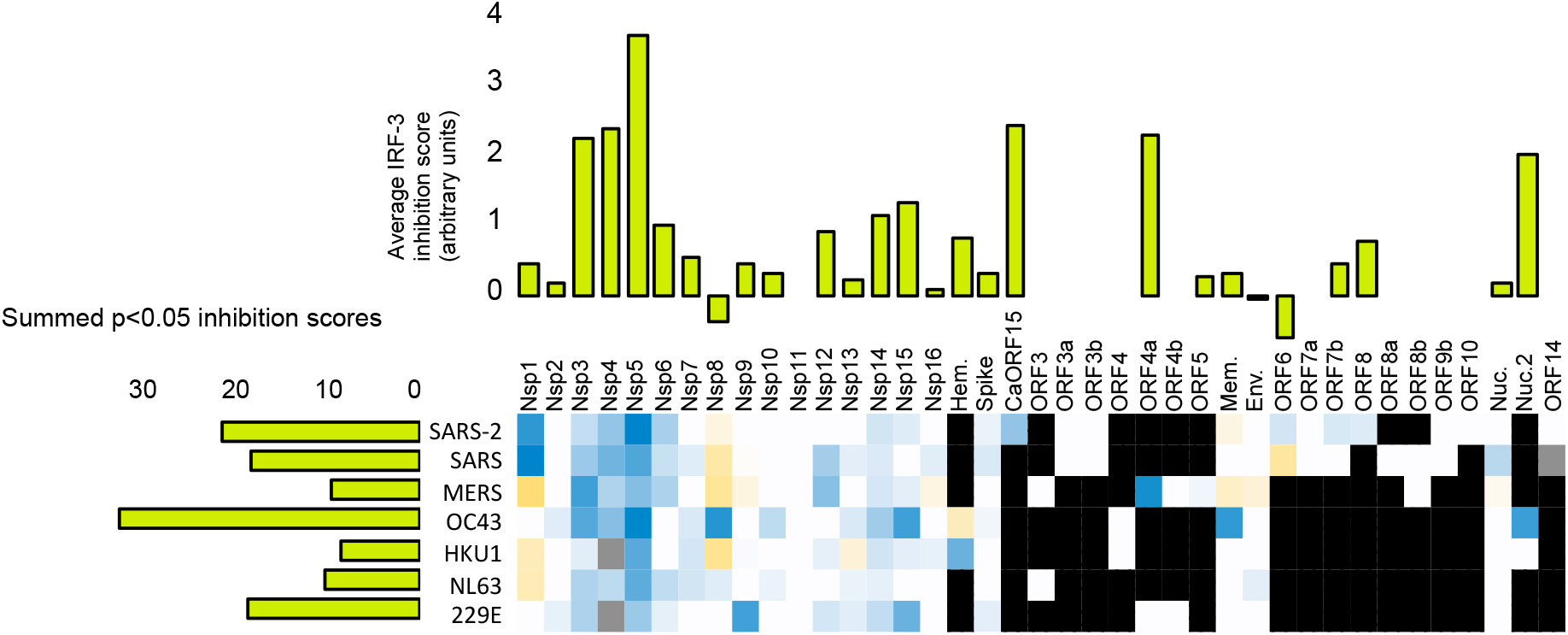
Inhibition of cGAMP activation of IRF-3 by coronavirus genes. This represents signaling through the STING pathway. These data, along with Supplementary Figures 2B and 2C are summed to generate Figure 3A.

**Supplementary Figure 2B.**
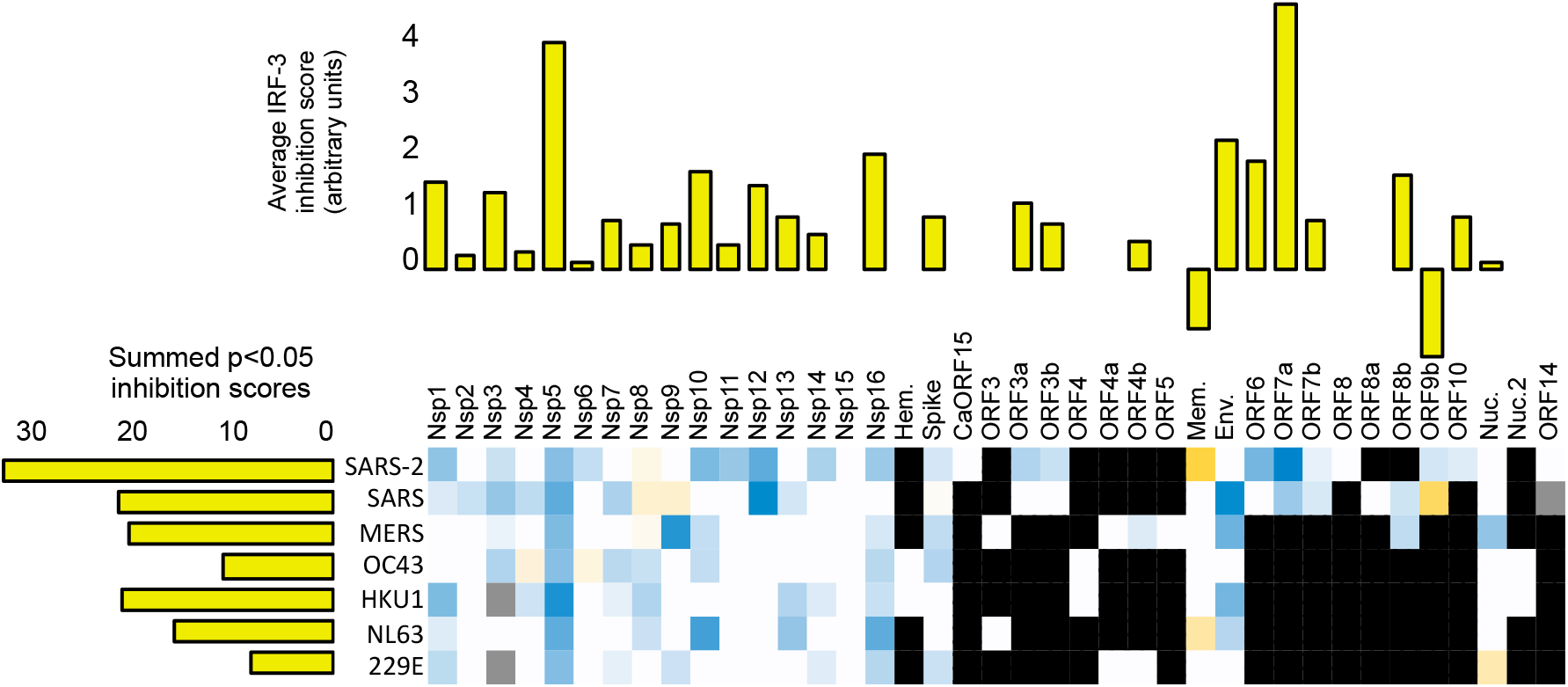
Inhibition of high-molecular weight polyI:C activation of IRF-3 by coronavirus genes. This represents signaling through the TLR-3 endosomal pathway.

**Supplementary Figure 2C.**
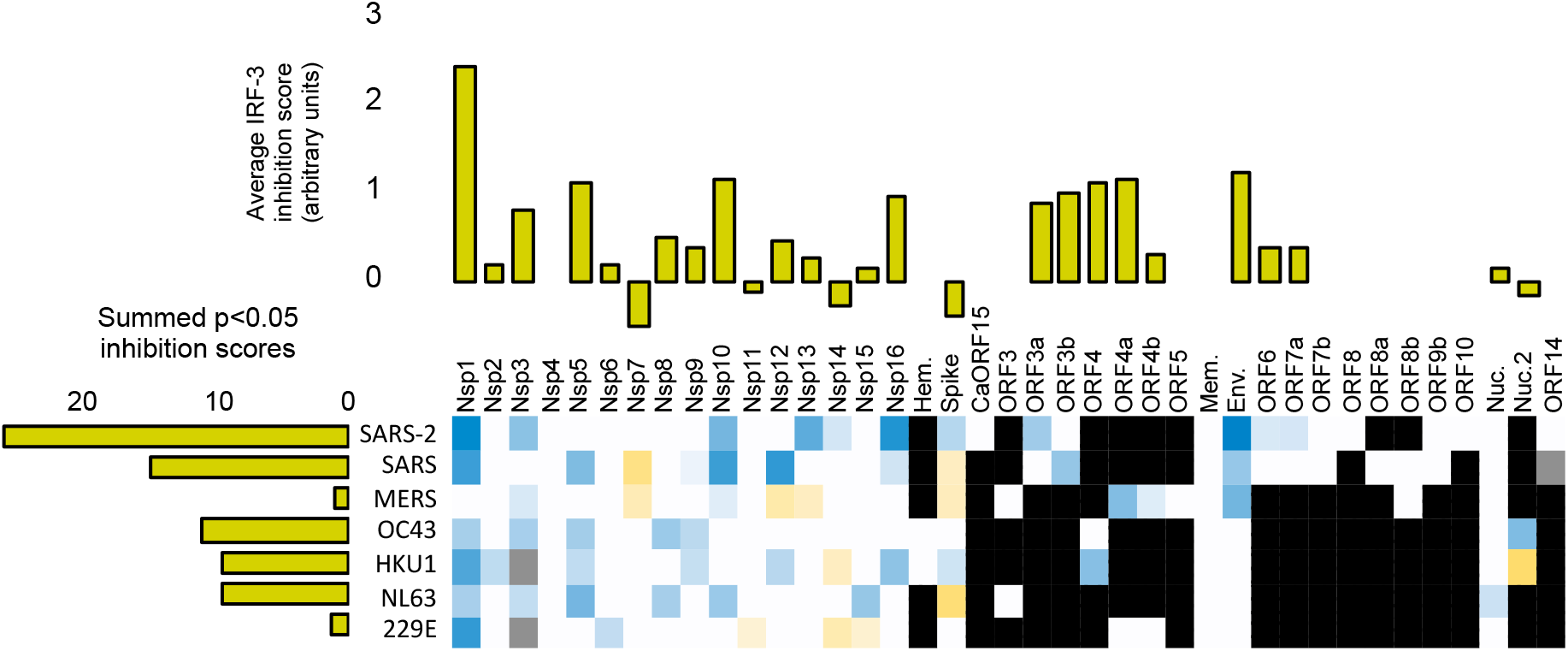
Inhibition of low molecular weight polyI:C/lipofectamine activation of IRF-3 by coronavirus genes. This represents signaling through the RIG-I-like receptor cytoplasmic pathway.

**Supplementary Figure 2D.**
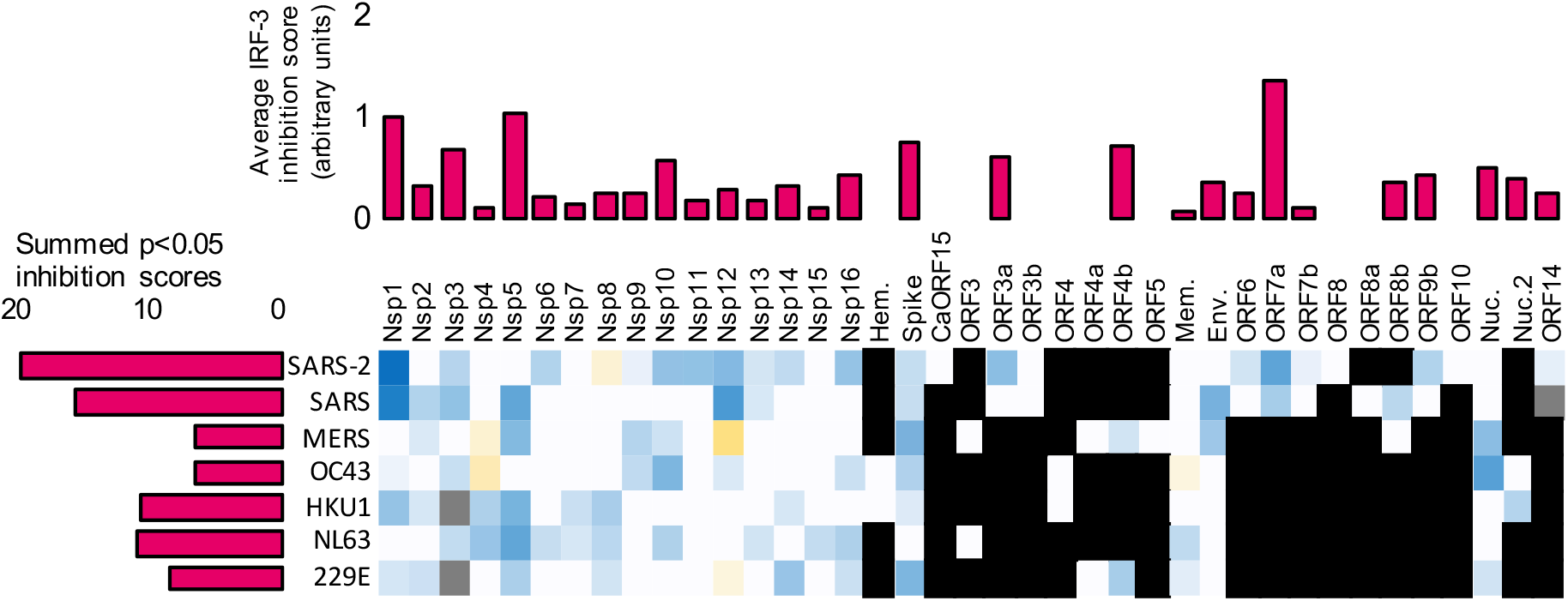
Inhibition of high-molecular weight polyI:C activation of NFkB by coronavirus genes. This presumably represents signaling through the TLR-3 endosomal pathway.

**Supplementary Figure 2E.**
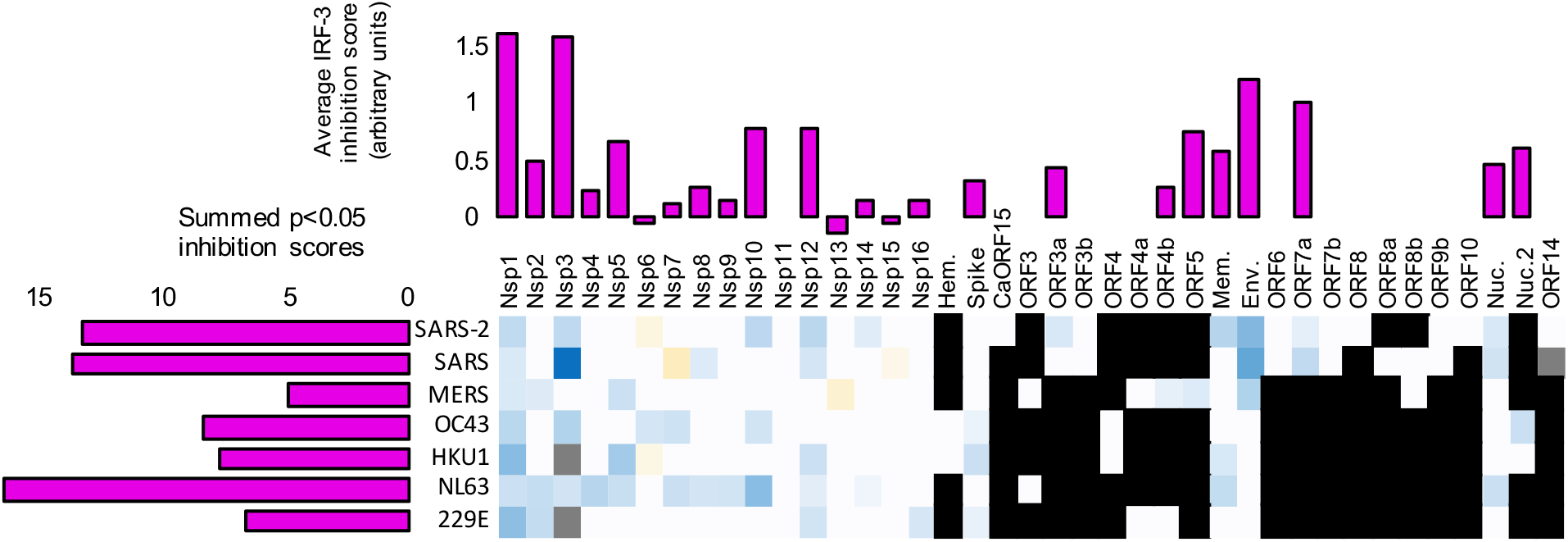
Inhibition of low molecular weight polyI:C/lipofectamine activation of IRF-3 by coronavirus genes. This presumably represents signaling through the RIG-I-like receptor cytoplasmic pathway.

**Supplementary Figure 2F.**
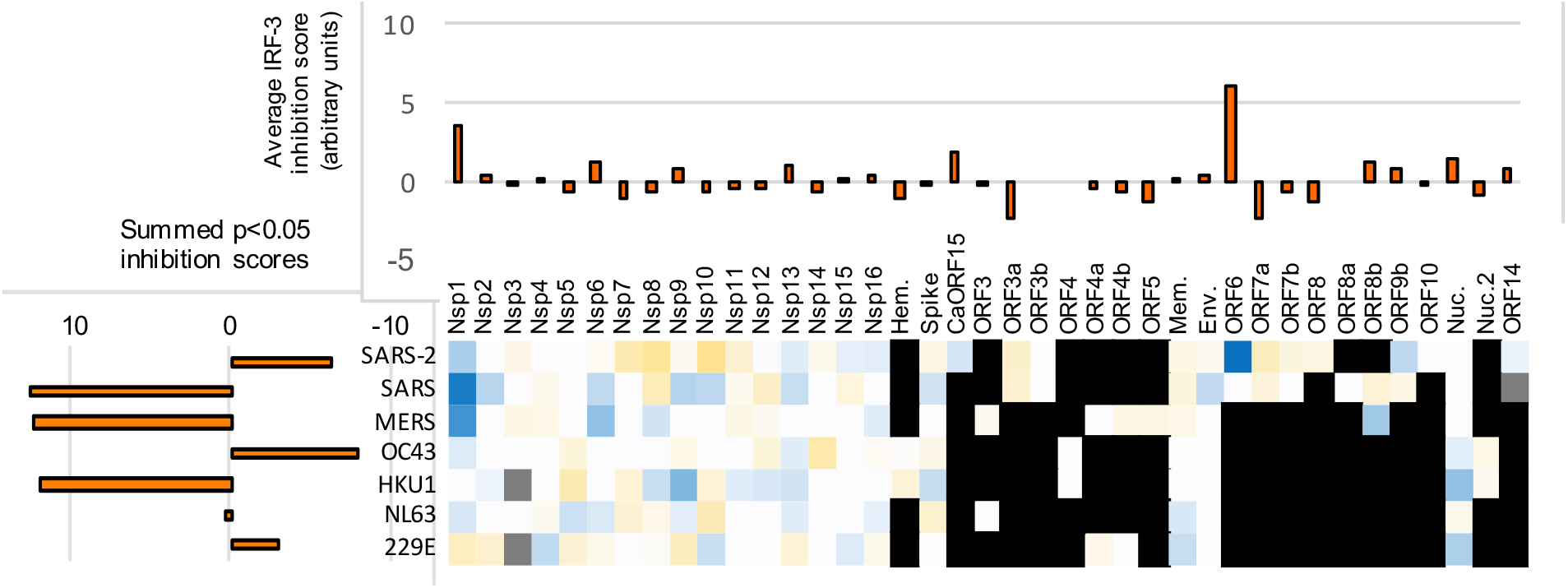
Inhibition by coronavirus genes of an increase in the nuclear/cytoplasmic ratio of phospho-STAT1 induced by interferon alpha.

**Supplementary Figure 2G.**
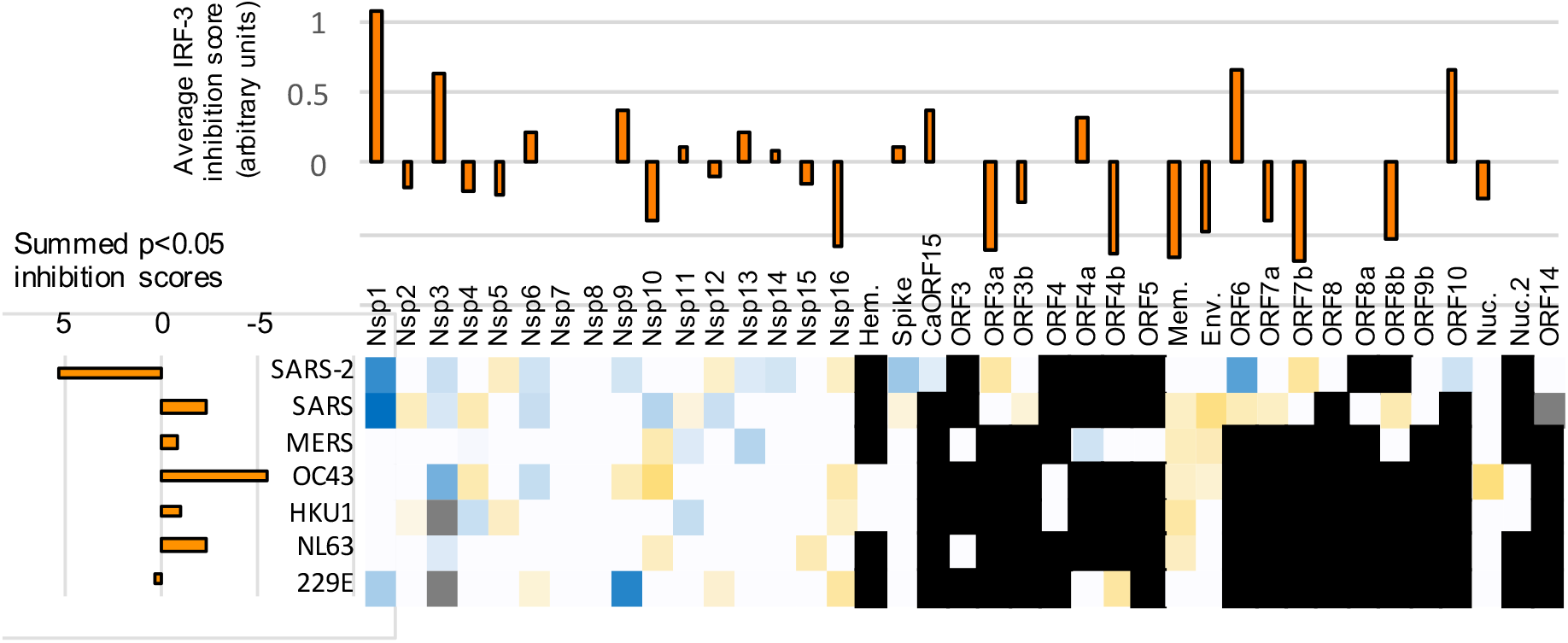
Inhibition by coronavirus genes of an increase in the nuclear/cytoplasmic ratio of STAT1 induced by interferon alpha.

**Supplementary Figure 2H.**
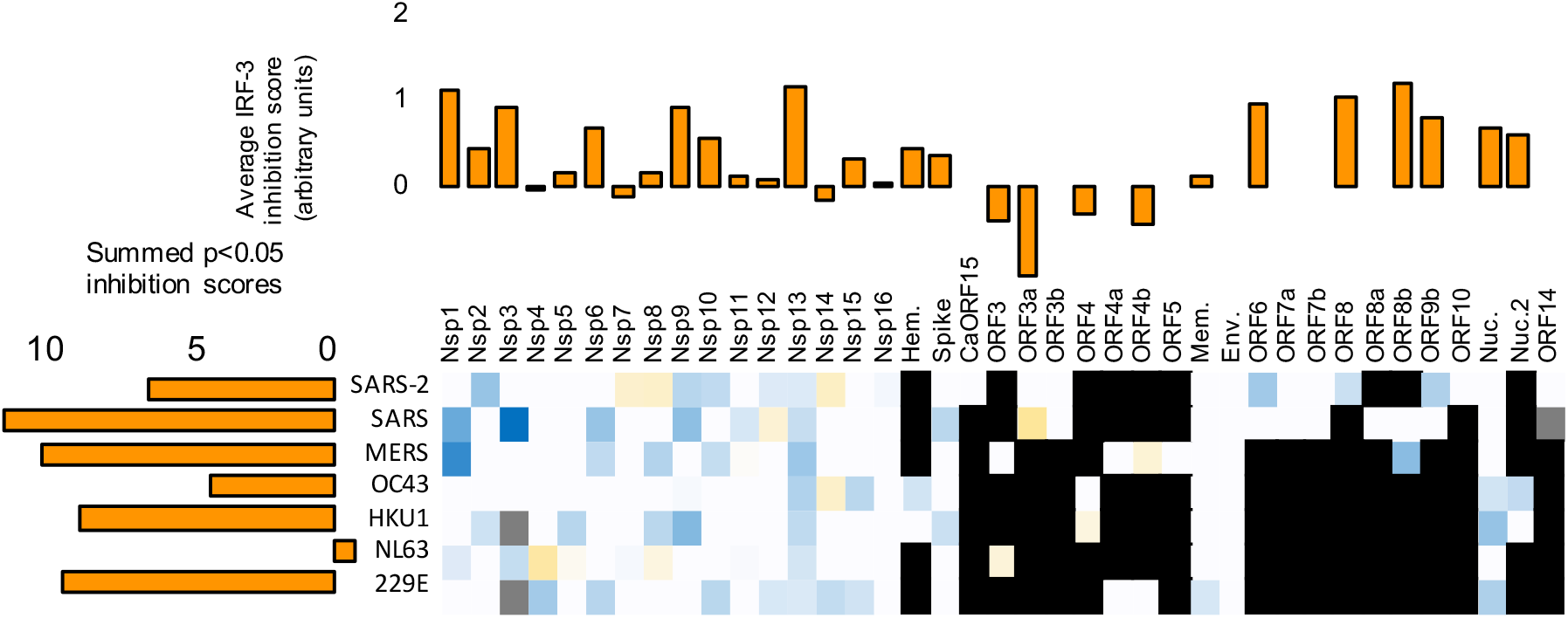
Effect of coronavirus genes on the total cellular amount of phospho-STAT1 induced by interferon alpha.

**Supplementary Figure 2I.**
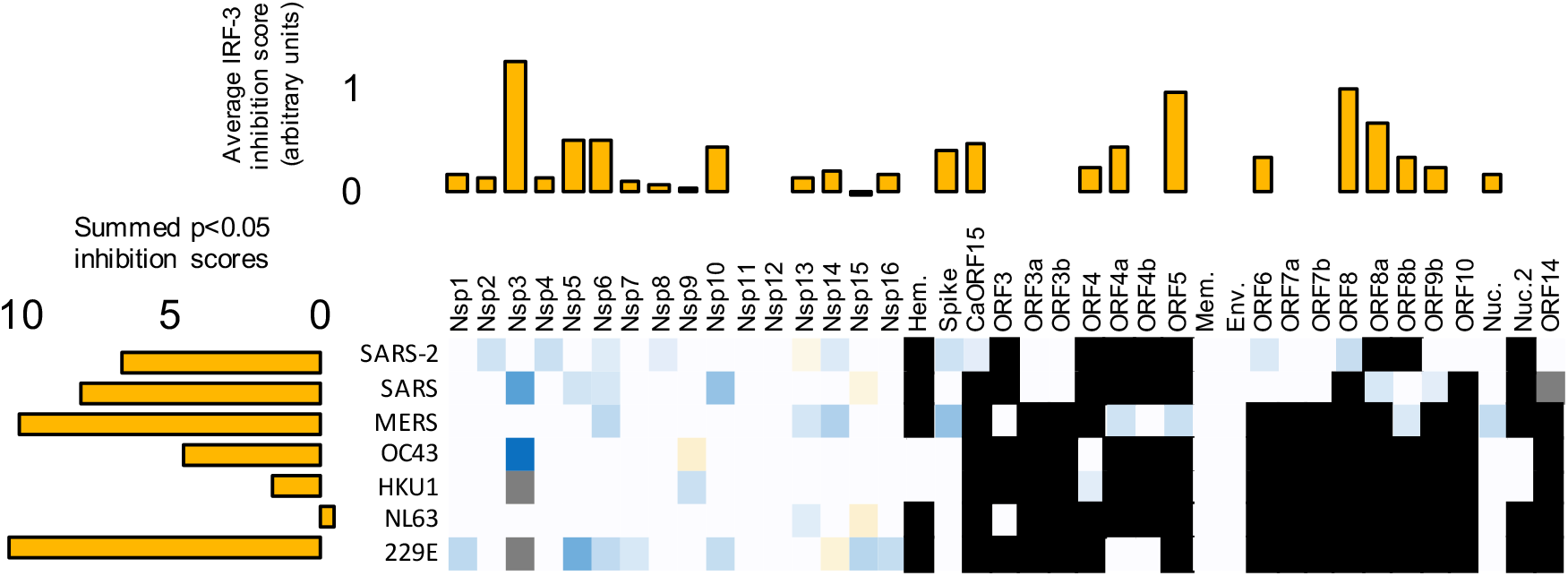
Effect of coronavirus genes on the total cellular amount of STAT1 after treatment by interferon alpha.

**Supplementary Figure 3.**
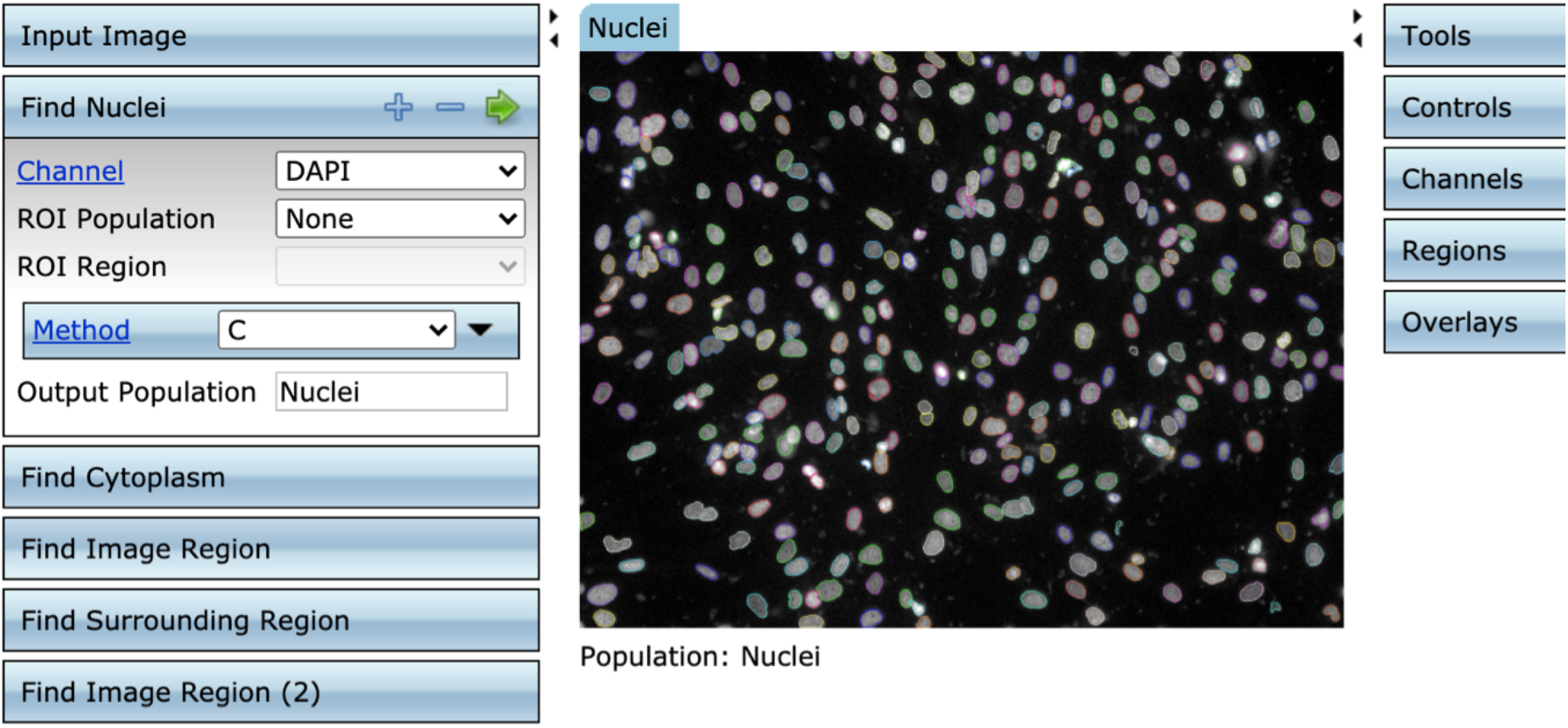
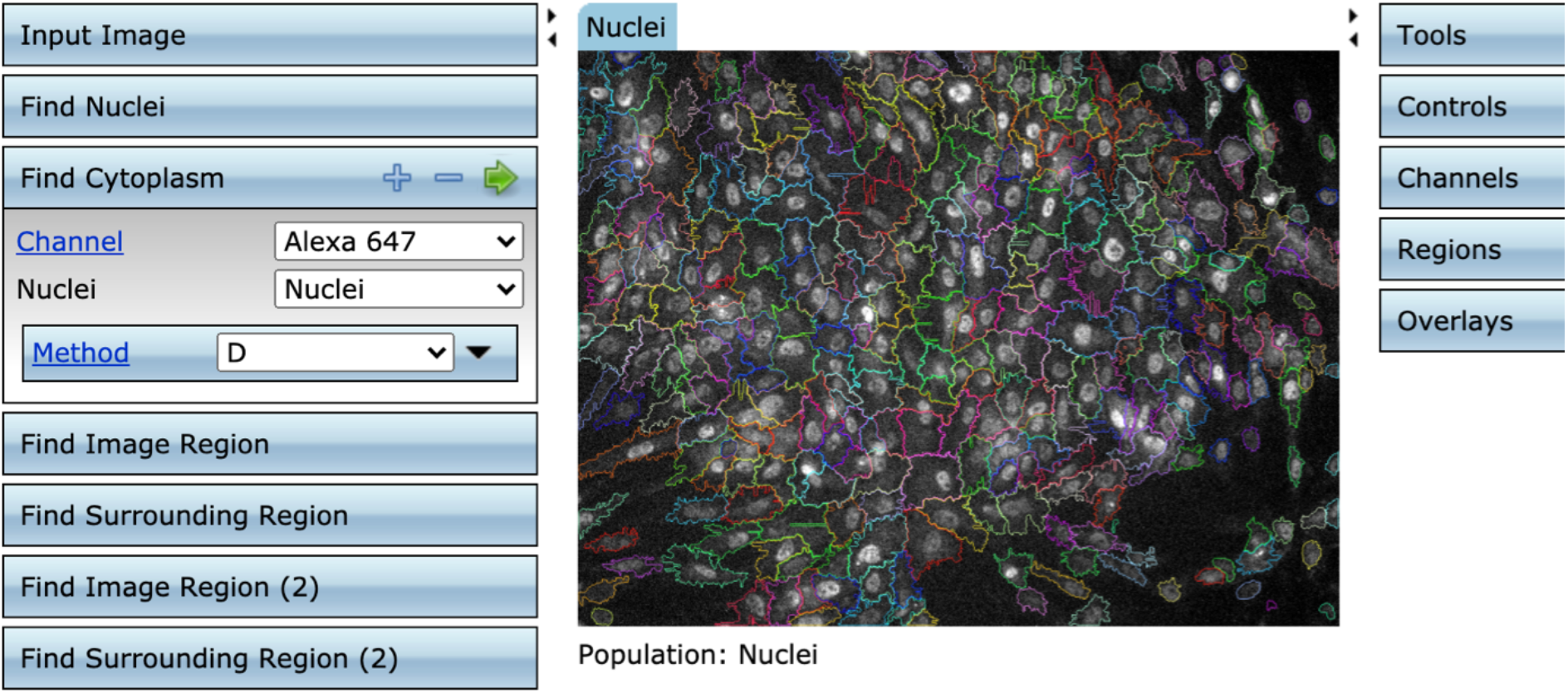
Image processing. Typical screen shot using the Columbus software, showing nuclear segmentation (top) and cytoplasmic segmentation (bottom).

**Supplementary figure 4.**
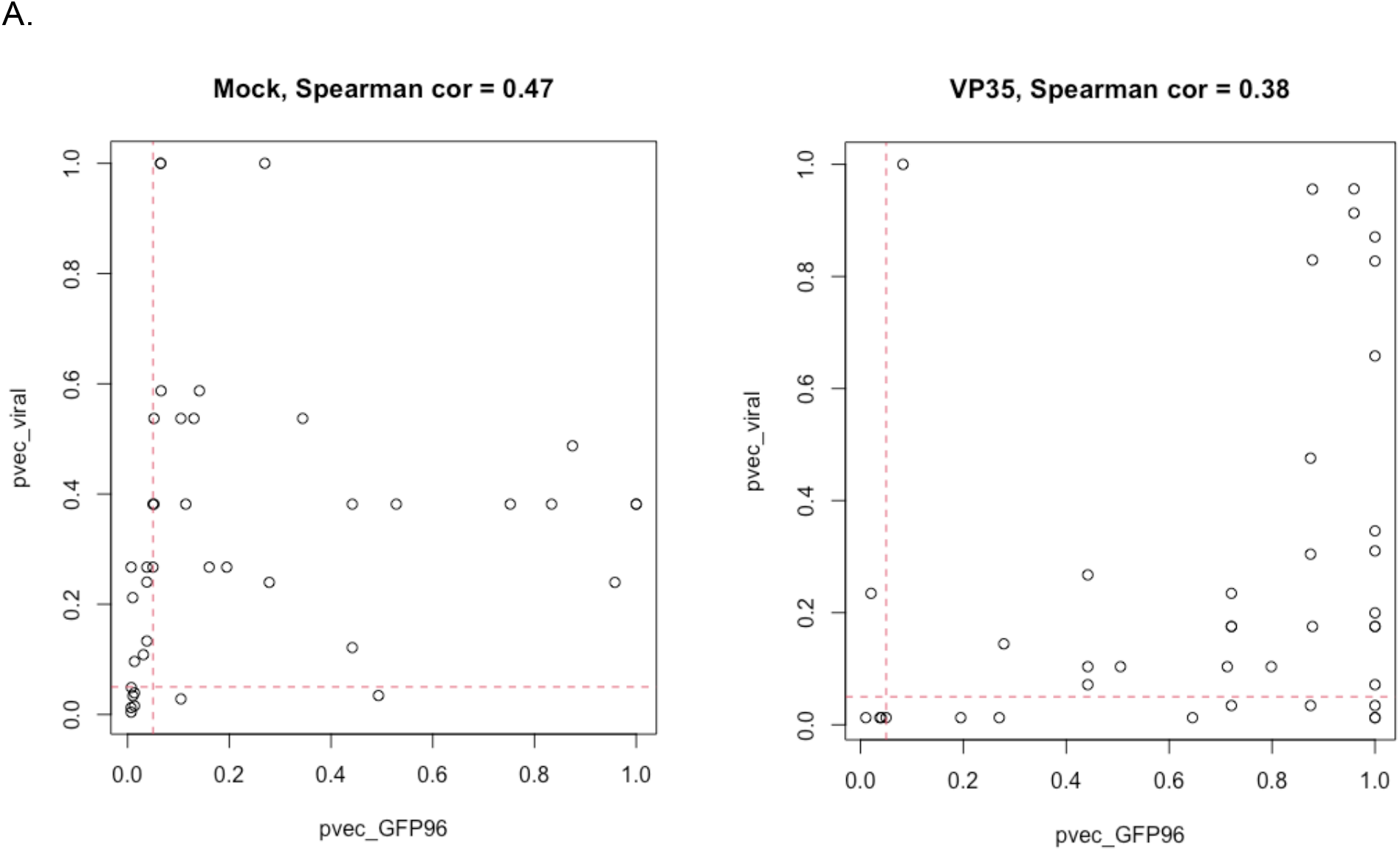
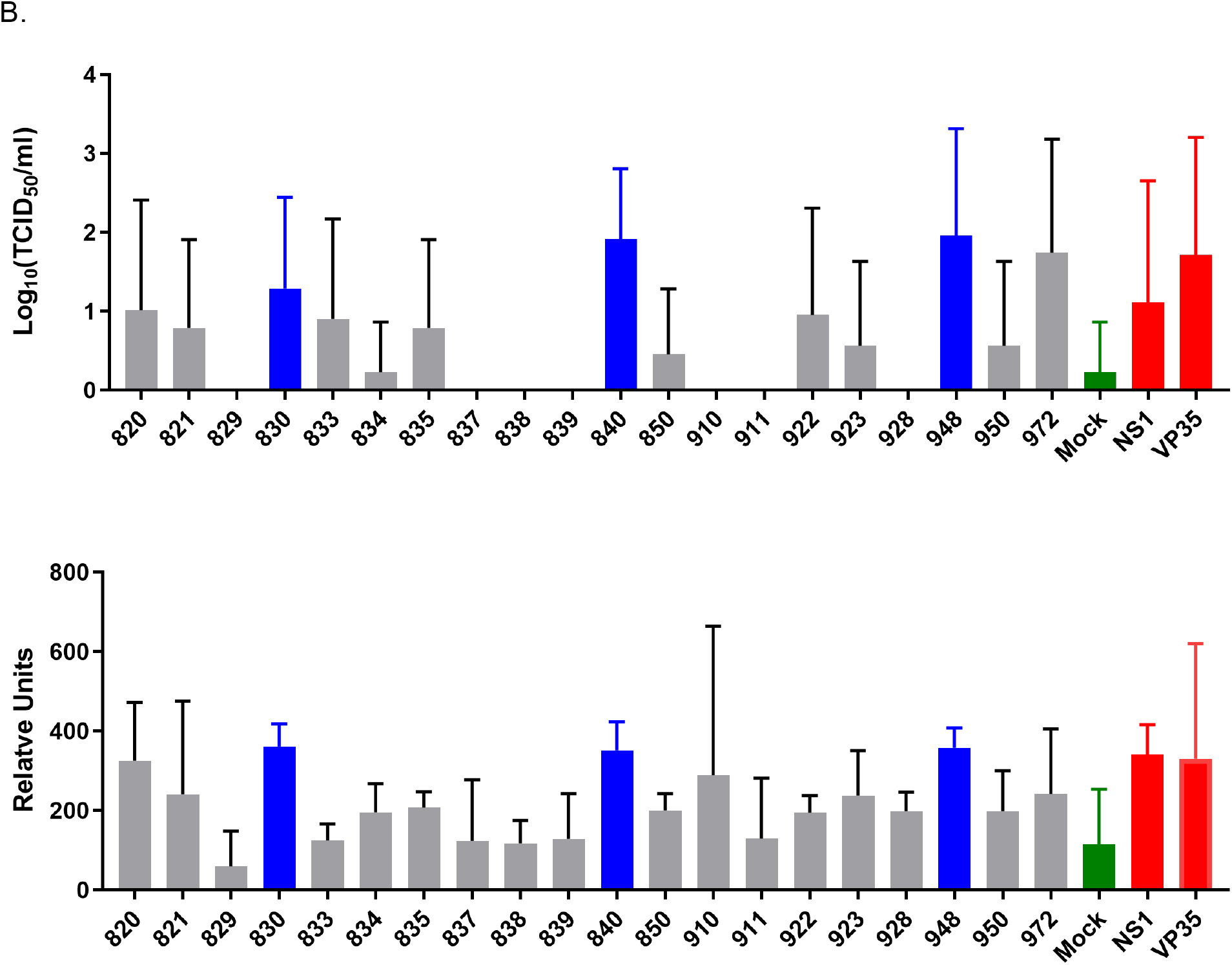
Comparison of HSV GFP relative fluorescence data to TCID_50_ viral replication data. A Wilcoxon rank sum test was used to assess the detection power of testing the difference between each coronavirus gene and positive/negative control using viral titers and GFP fluorescence with in the HSV-1 based assay. **A.** The negative (Mock) and positive (Influenza virus NS1 and Ebola VP35) control p-values were plotted on panel A and B, respectively. In each graph, x-axis is p-value using GFP fluorescence; y-axis is p-value using viral titers; red dashed line indicates p-value = 0.05. The hypothesis that if the two assay types produce similar results, then their p-values would have high correlation, suggesting similar detection power. Results showed a spearman correlation of 0.47 for mock (negative control) (panel A) and 0.38 for VP35 (positive control) (panel B), both indicating moderate correlation. **B.** The TCID_50_ viral replication titers from twenty coronavirus genes as well as the positive controls and negative control are shown in panel C and compared to the GFP data associated with each gene. Error bars indicate SD. Blue bars represent the coronavirus genes that were positive in the two assays and grey bars represent the coronavirus genes that were negative. Red bars are for the positive controls and the green bar is for the mock negative control.

## Notes

### Competing Interest Statement

The authors have declared no competing interest.

